# Topography influences diurnal and seasonal microclimate fluctuations in hilly terrain environments of coastal California

**DOI:** 10.1101/2023.09.22.559081

**Authors:** Aji John, Julian D. Olden, Meagan F. Oldfather, Matthew M Kling, David D. Ackerly

## Abstract

Understanding the topographic basis for microclimatic variation remains fundamental to predicting the site level effects of warming air temperatures. Quantifying diurnal fluctuation and seasonal extremes in relation to topography offer insight into the potential relationship between site level conditions and changes in regional climate. The present study investigated an annual understory temperature regime for 50 sites distributed across a topographically diverse area (>12 km^2^) comprised of mixed evergreen-deciduous woodland vegetation typical of California coastal ranges. We investigated the effect of topography and tree cover on site-to-site variation in near-surface temperatures using a combination of multiple linear regression and multivariate techniques. Sites in topographically depressed areas (e.g., valley bottoms) exhibited larger seasonal and diurnal variation. Elevation (at 10 m resolution) was found to be the primary driver of daily and seasonal variations, in addition to local topographic features that measure how depressed a site is compared to the surrounding area, canopy cover and northness. The elevation effect on seasonal mean temperatures was inverted, reflecting large-scale cold-air pooling in the study region, with elevated minimum and mean temperature at higher elevations. Additionally, several of our sites showed considerable buffering (dampened diurnal and seasonal temperature fluctuations) compared to average regional conditions measured by weather station. Results from this study help inform efforts to extrapolate temperature records across large landscapes and have the potential to improve our ecological understanding of fine-scale seasonal climate variation in coastal range environments.

## Introduction

Human-induced climate change is a major cause of species extinctions and biodiversity loss globally (Quintero and Wiens 2013). Widespread species range shifts in response to warming are already evident (Parmesan 2006, Chen et al. 2011), and substantial additional reshuffling of plant and animal communities is likely (Pecl et al. 2017). However, mounting evidence suggests that small scale topographic variation may promote high levels of climatic heterogeneity (Daly et al. 2009, Frey et al. 2016), thus modifying expected large scale range shifts caused by macroclimatic change (Parmesan 2006, Lenoir et al. 2013, Patsiou et al. 2014). Facilitated by unique micro-topographic characteristics (Dobrowski 2011, Patsiou et al. 2014), this can offer opportunities for species to persist in place, rather than shift in space, in response to climate change (Hampe and Petit 2005, Ashcroft et al. 2012, Thurman et al. 2020).

Topography and vegetation can create microclimates that differ from regional climates. Such departures can be explained by physio-topographic attributes such as elevation, slope and aspect, depressions and ridgetops, canopy cover, distance to coast, and proximity to the forest edge (Ewers and Banks-Leite 2013, Jucker et al. 2018). These features give rise to spatio-temporal heterogeneity expressed on local scales, affecting the magnitude of diurnal (daily maximum vs. minimum) and seasonal (summer vs. winter) temperature variations (Vanwalleghem and Meentemeyer 2009, Dobrowski 2011). Furthermore, the resulting minimum and maximum extreme temperatures, are known to structure plant and animal distributions (Sexton et al. 2009, Lembrechts et al. 2019). In mountain landscapes in particular, steep topographic gradients create strong microclimate heterogeneity.

Microclimatic buffering may manifest at diurnal, seasonal or interannual scales. We define it as reduced temperature fluctuations relative to free atmospheric conditions measured by e.g., weather station (Ashcroft et al. 2012, De Frenne et al. 2021). But, other definitions also exist; canopy resultant buffering of diurnal understory temperature fluctuations, with lower daytime maxima and higher nighttime minima (Greiser et al. 2018, Davis et al. 2018, De Frenne et al. 2019), and buffering of temperature extremes in comparison with a forest clearing e.g., clearcuts (Chen and Franklin 1997, De Frenne et al. 2013, Zellweger et al. 2019).

Cold-air pooling modifies the coupling of sites with free-air temperature in conjunction with physiographic features. Coupling is defined as having a 1:1 linear relationship with the free-air temperature (e.g., clearcut site or a weather station) (De Frenne et al. 2021). The cold-air pooling conditions keep cold air trapped in convergent environments like valley bottoms yielding stable microclimates in some circumstances (Schörghofer et al. 2018). Generally, the relationship between temperature and elevation is explained by the adiabatic lapse rate of 6-8°C decrease per 1000 meters increase in elevation. By contrast, the higher density of cold air can lead to accumulation in low-lying regions, and to temperature inversions with warm air resting above layers of cold air (Dorninger et al. 2011). This is predominantly manifested during the winter on clear nights with light winds. Hence, cold-air pooling is a key determinant of the degree of coupling between the boundary layer and free atmosphere (Lundquist and Cayan 2007, Burns and Chemel 2014).

Previous research has pointed to the role of micro-topographic and vegetative features in buffering or coupling of microclimates with respect to regional climate. However, what remains less clear is the way different landscape patterns (characterized by topographic features and vegetation) influence mean diurnal and seasonal temperature variations as such. In fact, the potential impacts of temperature variability and extremes can pose a greater risk to species than increases in mean temperature (Vasseur et al. 2014, Helmuth et al. 2014). It is likely that organisms occupying sites that have higher seasonal and diurnal fluctuations may exhibit greater tolerance of heat extremes and greater potential to withstand climate change impacts (Janzen 1967). This notion is not new: Janzen (1967) predicted decades ago that species experiencing large environmental variability will acclimate better and have increased range limits than the ones in lower variability regimes. Greater temperature fluctuations may prepare species better to tolerate future warming as they have evolved or acclimated to broader climatic tolerances.

Here, we study a large mixed-hardwood forest landscape in Northern California to determine the role of topography and vegetation in diurnal and seasonal fluctuations, with a focus on canopy cover, slope, aspect, and hillslope position. We specifically look at how topography drives diurnal and seasonal minimum, maximum, and mean temperatures, and influence resulting diurnal and seasonal temperature fluctuations. We hypothesize that valley bottom will show greater temperature fluctuations, due to the combination of cold-air pooling and reduced diurnal mixing with the local free atmosphere (Burns and Chemel 2014, Lareau and Horel 2015, Sheridan 2019), leading to reduced temperature buffering capacity compared to ridge-tops (conceptual model in Fig. 1).

**Figure 1:**
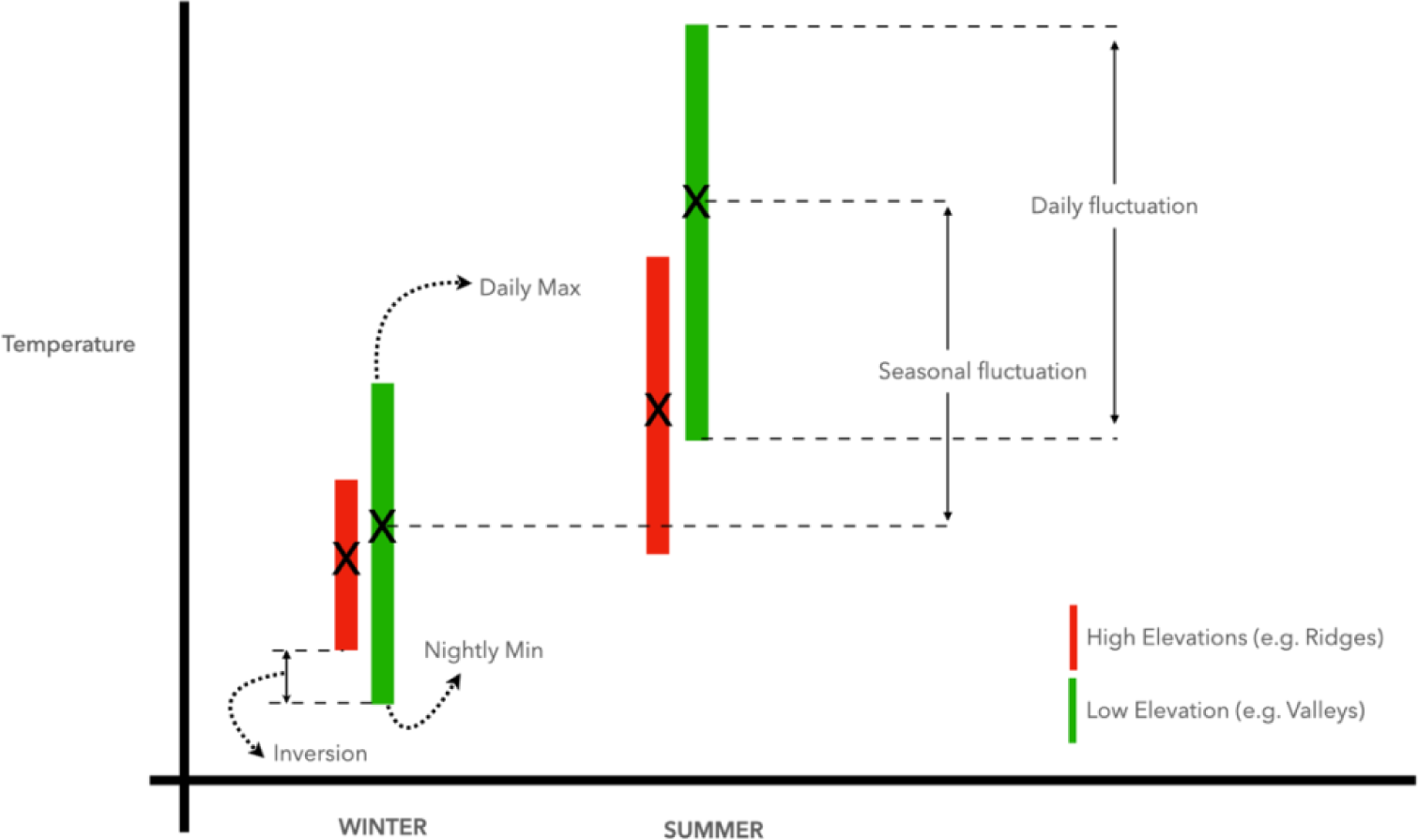
Conceptual diagram illustrating possible topographic effects on diurnal and seasonal fluctuations.

## Methods

### Study design

The study was conducted at Pepperwood Preserve (Sonoma Co., 38.57°N, -122.68°W) in northern California (Fig. 2: Pepperwood Preserve Study Area). The preserve is representative of deciduous and evergreen woodlands in the region, and exhibits elevations from 122 m to 462 m, with rugged undulating topography. Fifty 20 x 20 m vegetation plots were selected by stratifying on micro-topographic features and vegetation types (Oldfather et al. 2016).

**Figure 2:**
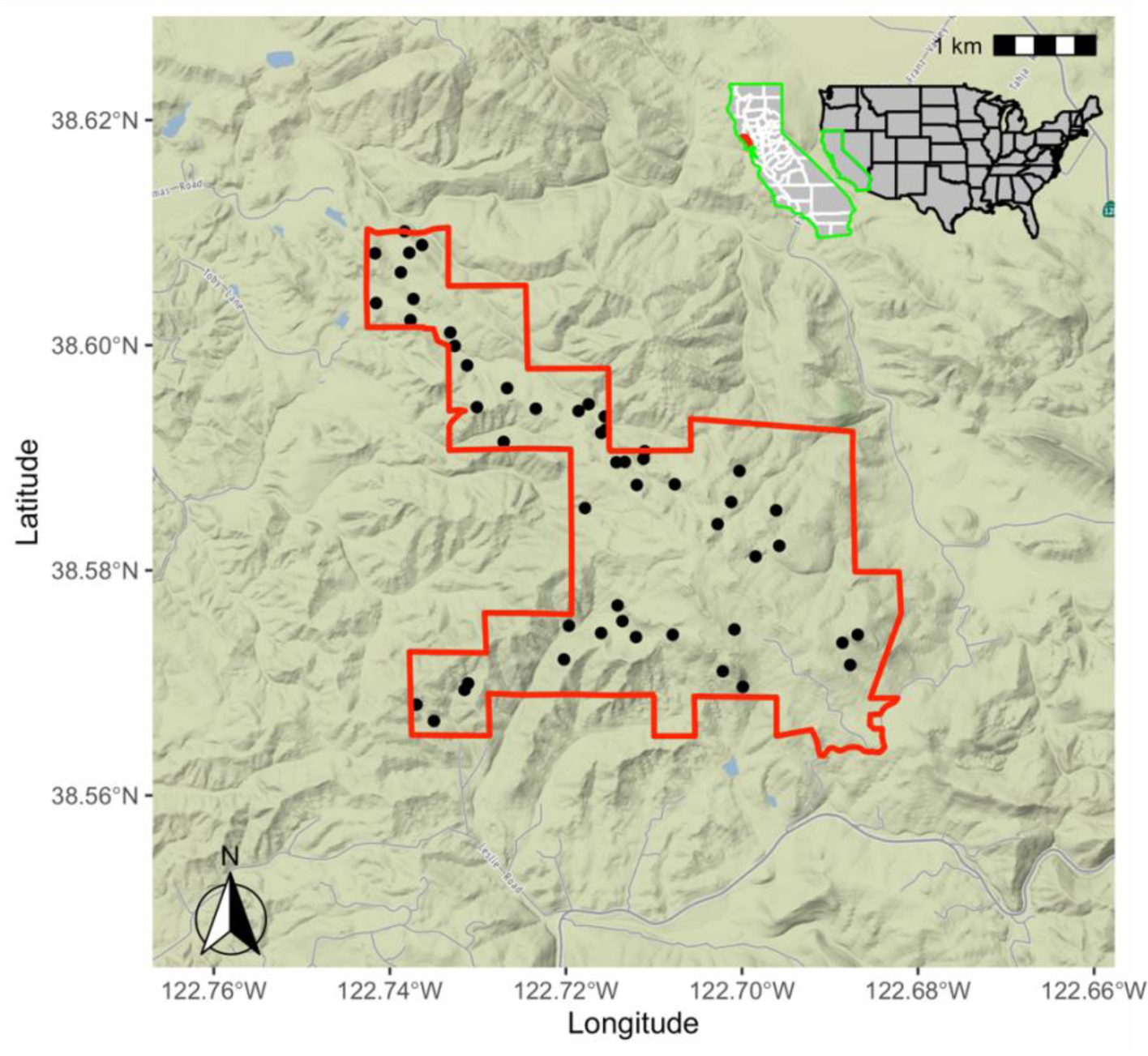
Pepperwood Preserve with 50 study sites. Each site was equipped with a microclimate logger (Onset HOBO Model U23, Onset Corp., Bourne, MA) that was installed at 1.2m height above the ground (see Fig. A.3 for the picture of the logger).

### Climate data collection

In the summer of 2013, 50 temperature data loggers (Onset HOBO Model U23, Onset Corp., Bourne, MA) were placed across the study area. Each logger was placed 1.2 m above the ground inside a radiation shield (Fig. A.3). Loggers were placed at the edge of each plot, and under representative canopy cover conditions. All of the loggers were placed in the understory and are part of a long-term forest dynamics research project (Oldfather et al. 2016, Ackerly et al. 2020). Data loggers recorded temperature (°C) and relative humidity (RH%) at half-hour intervals. At each site, average hourly temperatures were first calculated at the site level, and then from those average daily temperatures (minimum, maximum and diurnal) were calculated for winter (December 2013-February 2014), spring (March-May 2014), summer (June-August 2014) and autumn (Sep-Nov 2014) months. From these daily metrics, 18 summary statistics were then calculated (Table 1) representing daily minimum, mean, maximum, and diurnal variation in each season, plus growing degree day accumulation and the variation in seasonal means (difference between summer mean and winter mean). A permanent weather station at the Pepperwood Preserve was used as a reference in our study. To evaluate coupling and buffering of a site with regional (here weather station), quantification was done following methods of De Frenne et al. 2021.

**Table 1.**
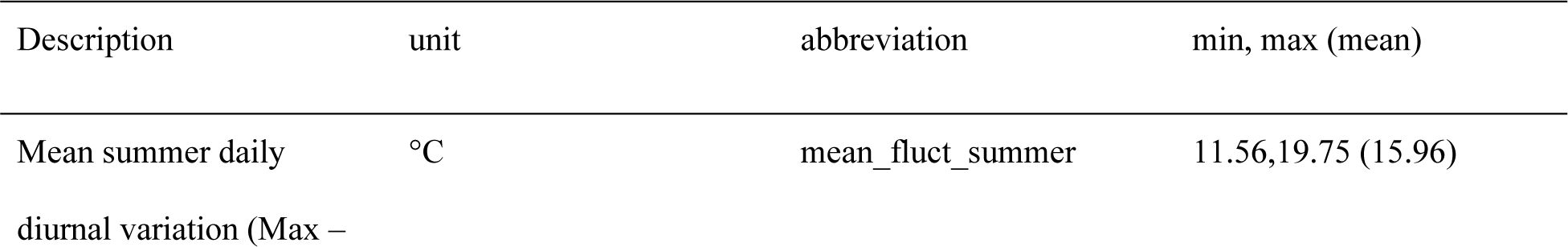
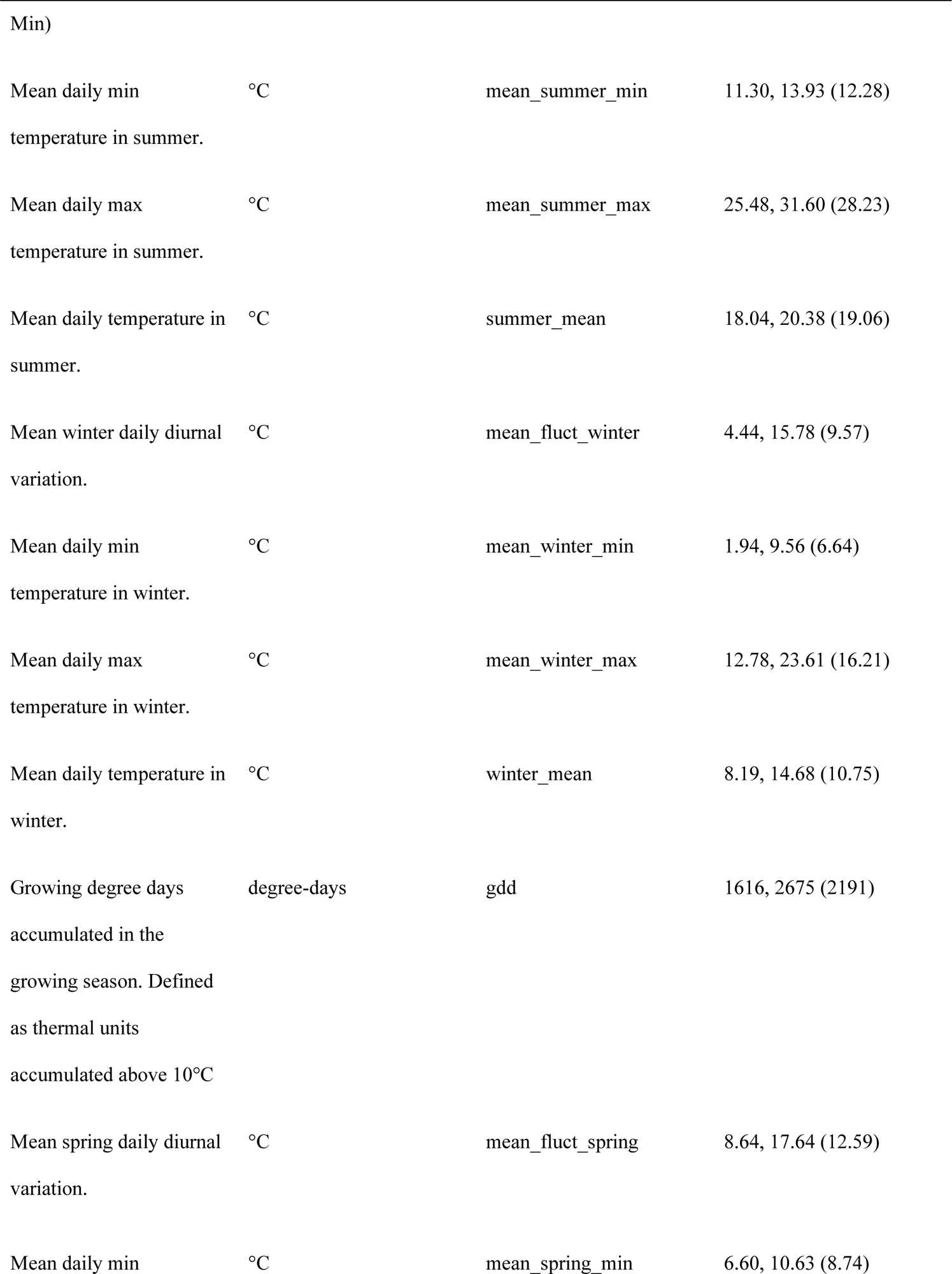
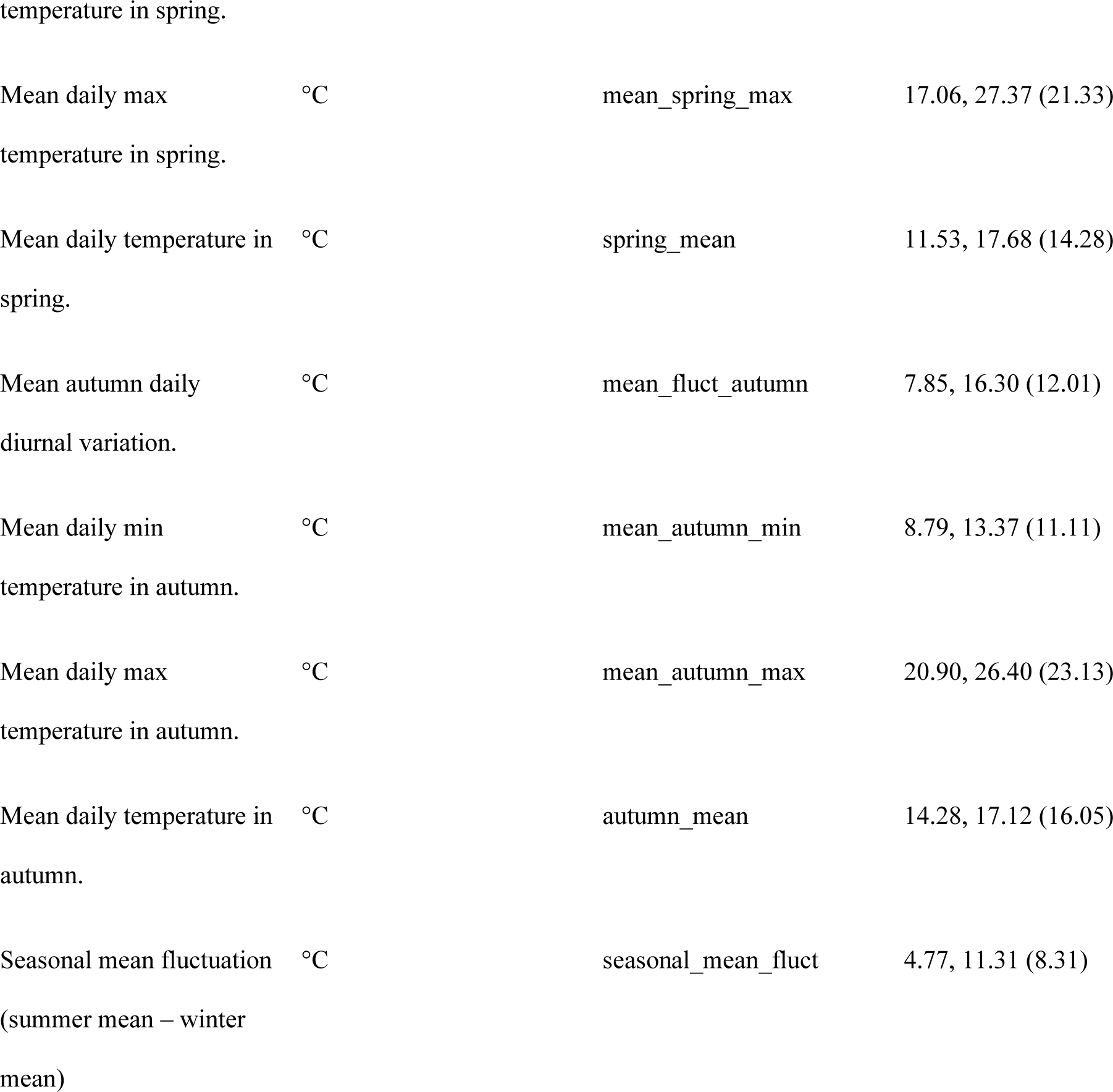
Derived climate variables and their minimum, mean, and maximum across the 50 logger sites.

### Topographic and canopy characterization

Spatial analysis using a GIS was performed on a 10 m digital elevation model (DEM) of Pepperwood Preserve to derive topographic variables (for details on topographic variables used in this study see Oldfather et al. 2016) (Table 2 and Table A.2). Variables included TPI (topographic position index), which is calculated as the difference (m) between the elevation of a point and the mean elevation in a surrounding radius. This measure indicates how elevated or depressed a site is in relation to its surroundings (positive for ridges and hilltops, and negative for valley bottoms). 100 m and 500 m radii were used to produce TPI100 and TPI500, respectively. TPI is essentially a local elevation, in contrast to DEM which is the global elevation. Similarly, PLP (percent lower points) is defined as the percentage of points within a given radius that are lower than the focal point (higher percentages indicate hilltops, and lower percentages identify valleys). PLP and TPI measures at each scale were strongly correlated (pairwise r^2^ ranged from .84 to .94), and PLP500 was selected for use in this analysis. Slope and aspect were calculated using the *raster* R package. Northness was calculated as cosine(aspect)*sine(slope) (Ackerly et al. 2020) (aspect on its own was examined but did not contribute to the final analysis). Pairwise Pearson correlation coefficients among topographic variables ranged from -0.26 to 0.29, with the highest between DEM and PLP500, as these are two measures of site elevation. Canopy cover was recorded in mid-summer when the plots were initially established, using a forest densiometer (Oldfather et al. 2016). Note that plots have varying proportions of deciduous trees (0 to 100%); the cover values are thus most relevant for analysis of summer temperatures. Canopy cover for winter was modified by subtracting the deciduous fraction.

**Table 2.** Topographic and canopy variables measured at each of the 50 sites.

| Description | unit | abbreviation | min, max (mean) |
| --- | --- | --- | --- |
| Elevation derived from Digital Elevation Model (referred as elevation in the text) | m | DEM (Elevation) | 121.89,461.52 (272.02) |
| Percent of pixels lower in a 500m radius | % | PLP500 (Topographic position) | 6.14, 99.85 (47.15) |
| Measure of steepness | ° | slope | 3.55, 29.77 (18.80) |
| Canopy cover | % | canopy | 32.85,91.03 (74.72) |
| Northness<br>($\cos(\text{aspect}) \cdot \sin(\text{slope})$) | | northness | -0.43, 0.42 (0.06) |

### Statistical analyses

Each site is a 20 m x 20 m plot and constitutes a study unit. Two data matrices were produced for analysis; the climate matrix contained all the climatic descriptors for the sites (50 sites x 18 variables), and the physiographic matrix included topographic and canopy related variables (50 sites x 5 variables). All the variables were continuous. Two sites (Site 1348 and 1350) were missing climate data for roughly six months, and therefore the inferred seasonal metrics were noted to be inaccurate but were still included in all the analyses for completeness. All analyses were performed using R software (R Core Team 2021).

To reduce the dimensionality and to assess dominant variables in climate and physiographic space, principal component analysis (PCA) was employed. PCA is known to work better than stepwise regression in multivariate datasets where high correlations exist between the variables. Correlation matrix was used for PCA as variables differed in their measurement units. PCA was performed separately on the physiographic and climate matrices. A Monte Carlo randomization test was used to assess the statistical significance of each principal component (PC) axis.

Multiple linear regression was performed to tease apart variations in climatic space for individual seasons. Additionally, principal component regression (PCR) was used to assess the association between physiographic characteristics (summarized according to PC1 and PC2) and the minimum, maximum, mean, and diurnal variations in summer and winter temperatures. A complementary redundancy analysis (RDA) was also conducted to investigate the relationship between the entire suite of climate and physiographic variables. We further performed selective RDA by subsetting seasonal (winter, summer, autumn and spring) climate variables and seeking out the dominant topographic variables contributing to corresponding diurnal variations in different seasons. Aggregate climate metrics (min, max, mean, and seasonal fluctuations) were normalized for RDA analysis using log-10 transformation.

## Results

Over the 12-month cycle (December 2013-November 2014), temperatures across the landscape varied from a minimum of -2.7°C to a maximum of 40.4°C; mean hourly temperatures across the seasons (over all the days in a season) were between 9°C and 30°C (Fig. 3A-C), with an overall site mean temperature of 15°C (over all sites and days). Sites across the Pepperwood Preserve show significant diurnal variations in different seasons. In summer, the average diurnal variation was approximately 16°C and in winter it was approximately 9°C. Winter maximum temperatures exhibited the greatest spatial variability across sites (Fig. 3D).

**Figure 3:**
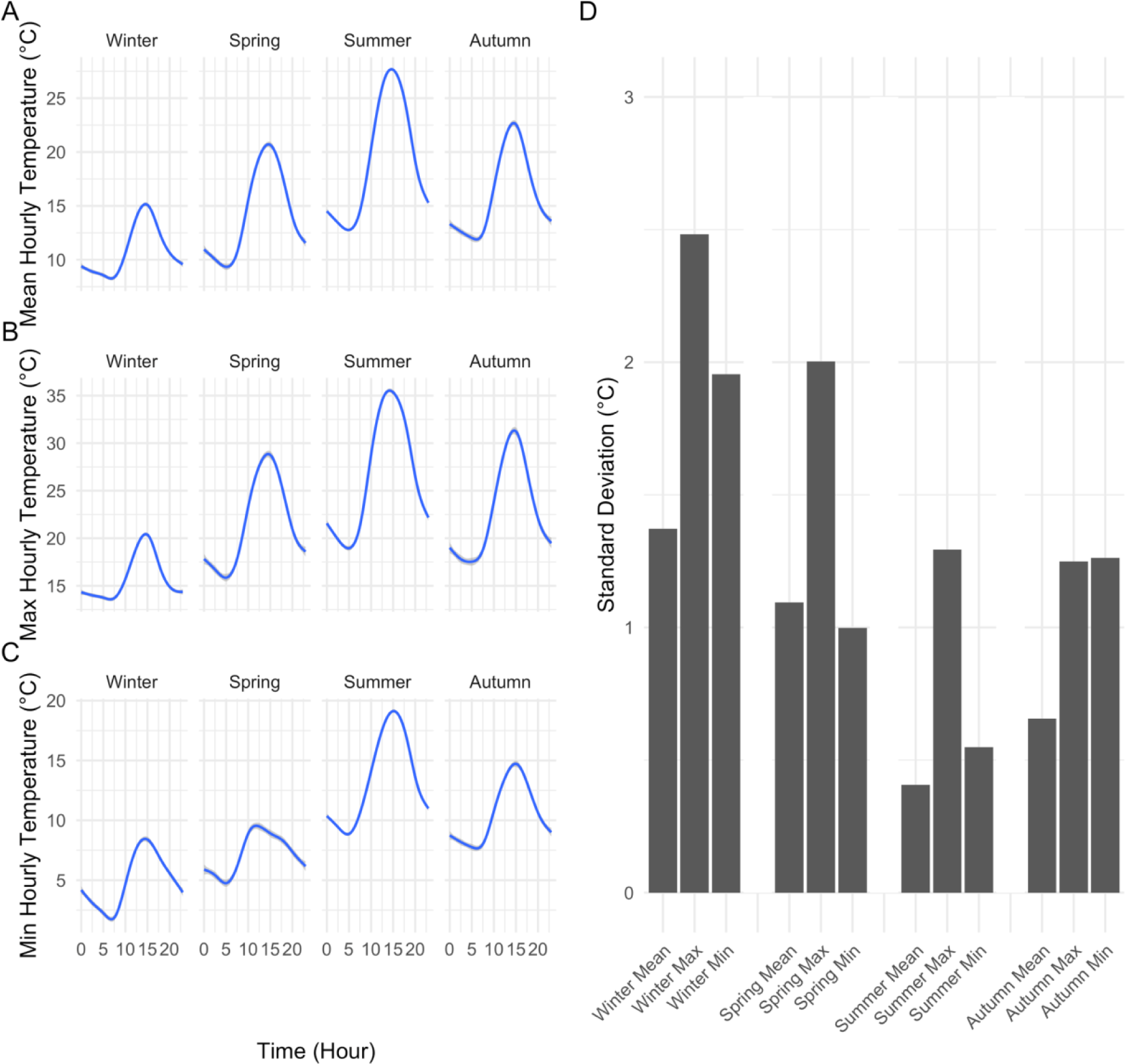
Seasonal minimum, maximum average temperatures with corresponding variability (lines are loess fits). (a) Mean hourly temperatures of 50 sites by season (b) Maximum hourly temperatures of 50 sites by season (c) Minimum hourly temperatures of 50 sites by season (d) Standard deviation of seasonal mean, minimum and maxima across sites.

Multiple regression analysis revealed that seasonal climate metrics were strongly related to elevation and moderated by micro-topographic and canopy features (Table 3). Diurnal variations in each season decreased with DEM (elevation), indicating that higher elevations exhibited lower diurnal variations than did lower elevation sites. Diurnal variations for autumn, spring and summer (all seasons except winter) were correlated negatively with PLP500 and canopy cover. Diurnal variation was also found to be negatively associated with northness in autumn only. Seasonal minimum temperatures were positively associated with elevation, suggesting temperature inversions, in contrast with max temperatures that were negatively associated with elevation. The positive association of minimum temperatures were primarily with DEM (elevation) and a finer elevation measure of PLP500 except in winter where it was elevation only. Seasonal maximum temperatures were negatively associated with elevation but were found to be moderated by canopy cover in summer and in autumn. Additionally, autumn and winter maximum temperatures were negatively associated with northness. Seasonal fluctuations (summer mean minus winter mean) decreased with elevation only.

**Table 3.** Partial coefficients from the multiple linear regression of climate variables by topographic drivers (*p-value* significance codes in bold: ‘***’ < 0.001, ‘**’ < 0.01, ‘*’ < 0.05, ‘.’ < 0.1, ‘’ < 1).

| Variable | Beta coefficients of main physiographic variables |  |  |  |  | R-Squared |
| --- | --- | --- | --- | --- | --- | --- |
|  | DEM | Slope | Canopy | PLP500 | Northness |  |
| mean_fluct_winter | **-0.018729\*\*\*** | 0.046583 | -0.018464 | -0.016301 | -3.283823 | 0.42 |
| mean_fluct_spring | **-0.015305\*\*\*** | -0.025467 | **-0.036223\*** | **-0.025515\*\*** | -0.450737 | 0.68 |
| mean_fluct_summer | **-0.010699\*\*\*** | 0.002919 | **-0.049029\*\*\*** | **-0.013381\*** | 0.265929 | 0.72 |
| mean_fluct_autumn | **-0.016498\*\*\*** | -0.006297 | **-0.040870\*\*\*** | **-0.021807\*\*\*** | **-1.780846\*\*** | 0.86 |
| winter_mean | **0.007027\*\*** | 0.041013 | -0.005661 | 0.008674 | -1.291933 | 0.34 |
| spring_mean | -0.002730 | -0.022272 | -0.012030 | 0.009555 | -0.013236 | 0.11 |
| summer_mean | -0.000377 | -0.008442 | **-0.020732\*\*\*** | **0.004923\*** | 0.097691 | 0.59 |
| autumn_mean | **0.003167\*\*\*** | -0.004384 | **-0.013655\*\*** | **0.013543\*\*\*** | **-1.041096\*\*\*** | 0.64 |
| winter_min | **0.015590\*\*\*** | 0.026200 | 0.000031 | 0.013930 | -0.504000 | 0.62 |
| spring_min | **0.004044\*\*** | -0.011007 | 0.002343 | **0.021983\*\*\*** | -0.287895 | 0.51 |
| summer_min | **0.002890\*\*\*** | -0.004501 | 0.001565 | **0.012609\*\*\*** | -0.201848 | 0.66 |
| autumn_min | **0.009473\*\*\*** | -0.003004 | 0.001554 | **0.022527\*\*\*** | -0.716712 | 0.80 |
| winter_max | -0.003137 | 0.072785 | -0.018495 | -0.002374 | **-3.787788\*** | 0.21 |
| spring_max | **-0.011261\*\*\*** | -0.036474 | -0.033880 | -0.003532 | -0.738632 | 0.36 |
| summer_max | **-0.007807\*\*\*** | -0.001582 | **-0.047463\*\*\*** | -0.000771 | 0.064080 | 0.66 |
| autumn_max | **-0.007025\*\*\*** | -0.009300 | **-0.039316\*\*\*** | 0.000720 | **-2.497557 \*\*\*** | 0.67 |
| seasonal_amplitude | **-0.007396 \*\*\*** | -0.051402 | -0.016380 | -0.002657 | 1.143957 | 0.38 |
| (summer and<br>winter) |  |  |  |  |  |  |
| gdd | -0.224700 | -4.975600 | **-8.044800\*\*\*** | **3.328300\*\*** | **-375.866000\*\*** | 0.44 |

Topographically, strong negative elevation effects were found in diurnal variations for all the seasons. PLP500 (direct measure of hilltop/valley bottoms) was also found to be negatively correlated with diurnal variation except in winter, i.e., hilltops/ridges show lower diurnal variations compared to valley bottoms. Canopy cover was found to be negatively associated with diurnal variations except in winter season and was found to influence summer and autumn mean and max temperatures. In addition, northness was found to influence autumn metrics (diurnal fluctuation, mean and max temperature), and winter max temperatures; additionally, it was found to influence growing degree days. Slope was not found to play any role on any of the variables.

Significant proportions of the variability were captured by the first two PCs in climate and physiographic space. In the first PCA (Climate), PC1 and PC2 explained 37.2% and 30.0%, respectively, of the climatic variability across Pepperwood Preserve (Fig. 4A). Variables with positive loadings on PC1 included mean minimum temperatures from winter, summer, and autumn, and winter and autumn mean temperatures. Variables with negative loadings on PC1 included summer and winter fluctuations and seasonal maxima. Positive loadings on PC2 included GDD, and several seasonal means and maxima. In the ensuing PCA results, we show the R^2^ of the respective correlations in parentheses. In the second PCA (physiographic space), PC1 and PC2 explained 54.3 % (27.2% and 27.1% respectively) of the topographic variability across Pepperwood Preserve (Fig. 4B). Topography related variables were positively correlated with PC1 except for DEM and PLP500 which was negatively correlated: DEM (R^2^ = 77%), slope (R^2^ =42%), northness (R^2^ = 42%) and PLP 500 (R^2^ = 7%). Canopy was positively correlated on PC1 (R^2^ = 74%). Hence, positive values of PC1 represent sites that are high elevation north facing ridges with steep slopes and denser canopy. PC2 of physiographic space was largely composed of positively correlated topographic variables: DEM (R^2^ = 35%), PLP 500 (R^2^ = 88%) and northness (R^2^ = 63%). Canopy and slope were positively correlated on PC2 (R^2^ = 20% & 14%). Positive values of PC2 can be interpreted as low canopy cover sites on higher elevations (ridgetops).

**Figure 4:**
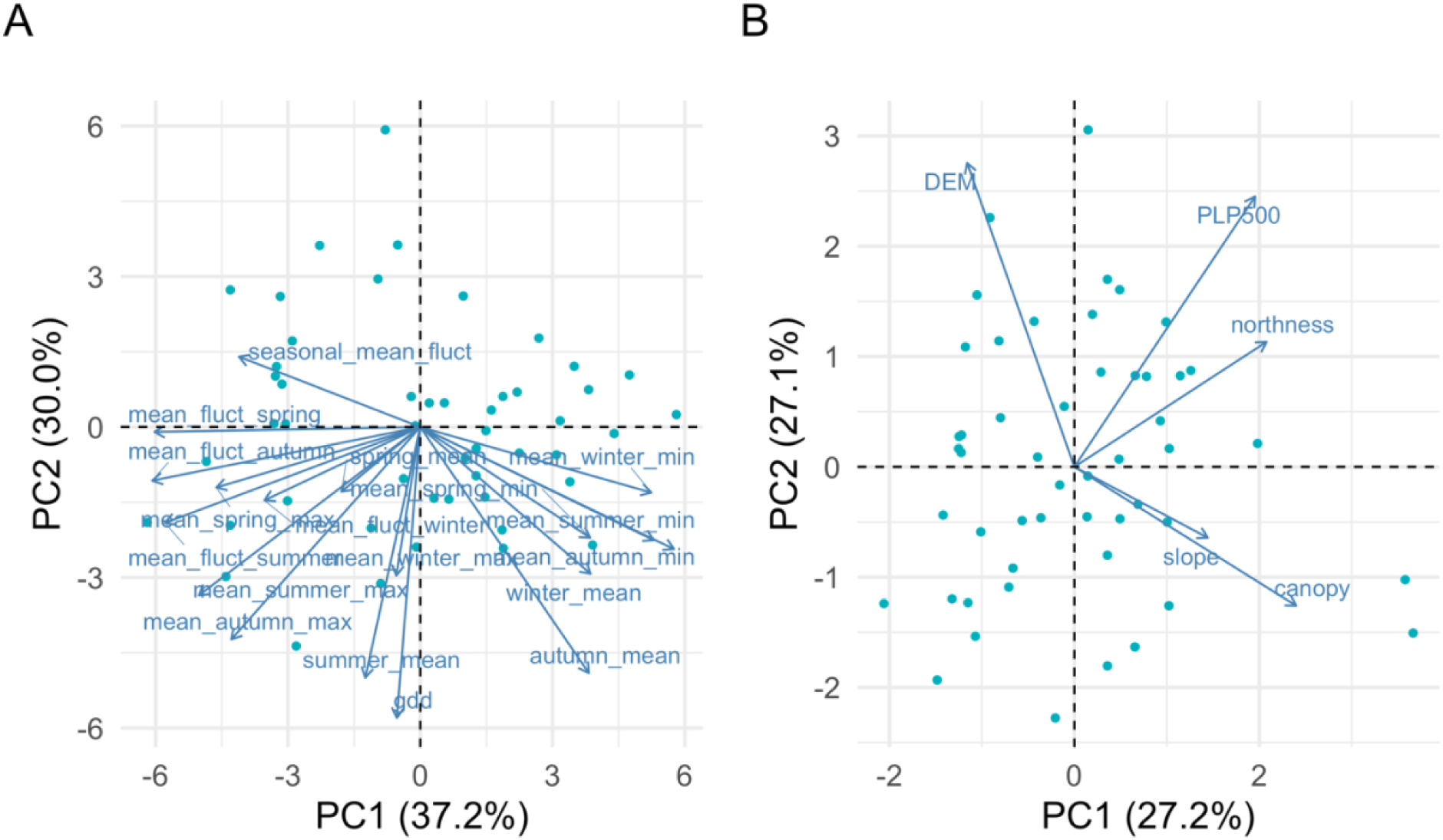
Biplots of climate and physiographic PCA analysis (a) PCA of climate space (b) PCA of physiographic space. Sites (points) are referenced to the lower and left axes scales, and variables (blue arrows) are referenced to the upper and right axes scales.

A multivariate linear regression model with the physiographic PCs was explored against all the climate variables (Fig. A.1, Fig. A.2). Mean summer and summer maxima were positively correlated with PC1 of physiographic PCA axes (R^2^ = 37% & R^2^ = 23%, p-value < 0.0001), summer diurnal variations were marginally correlated (R^2^ = 13%, p-value < 0.05), however, winter temperatures measures were found to be not correlated with PC1. PC2 of topographic space reflected the gradient from valley bottoms to high elevation sites and explained variation in winter temperature measures. Winter minimum and winter mean temperatures were positively correlated with PC2 of physiographic PCA axes (R^2^ = 42% & R^2^ = 9%, p-value < 0.0001 and p-value < 0.05), but winter diurnal variations and maximum temperatures were negatively correlated (R^2^ = 37% & R^2^ = 7%, p-value < 0.0001 and p-value < 0.05) with PC2 of physiographic PCA axes (Fig. A.2).

Redundancy analysis (RDA) confirmed the strong influence of topographic variables contributing to climate variability. RDA results show that topographic variables DEM, PLP500, northness and slope explained a significant proportion of climatic variability across all the seasons (Fig. 5). Table A.1 shows the RDA output when all the topographic variables were selectively constrained against seasonal climate variables. In autumn, topographic features (DEM, PLP500, northness) and canopy are found to be statistically significant (p-value < 0.005), but in spring, topographic features DEM and northness are significant (p-value < 0.005). In summer, DEM, PLP500 and canopy are found to be statistically significant (p-value < 0.005), and in winter, only DEM is found to be statistically significant (p-value < 0.005). Additionally, temperature metrics (maximum and diurnal) overlap with canopy and slope explaining the seasonal variation in all seasons, but canopy plays an important role in summer (Fig. 5). RDA analysis shows that DEM is the most dominant topographic variable which explains the variation in climate across a heterogeneous landscape.

**Figure 5:**
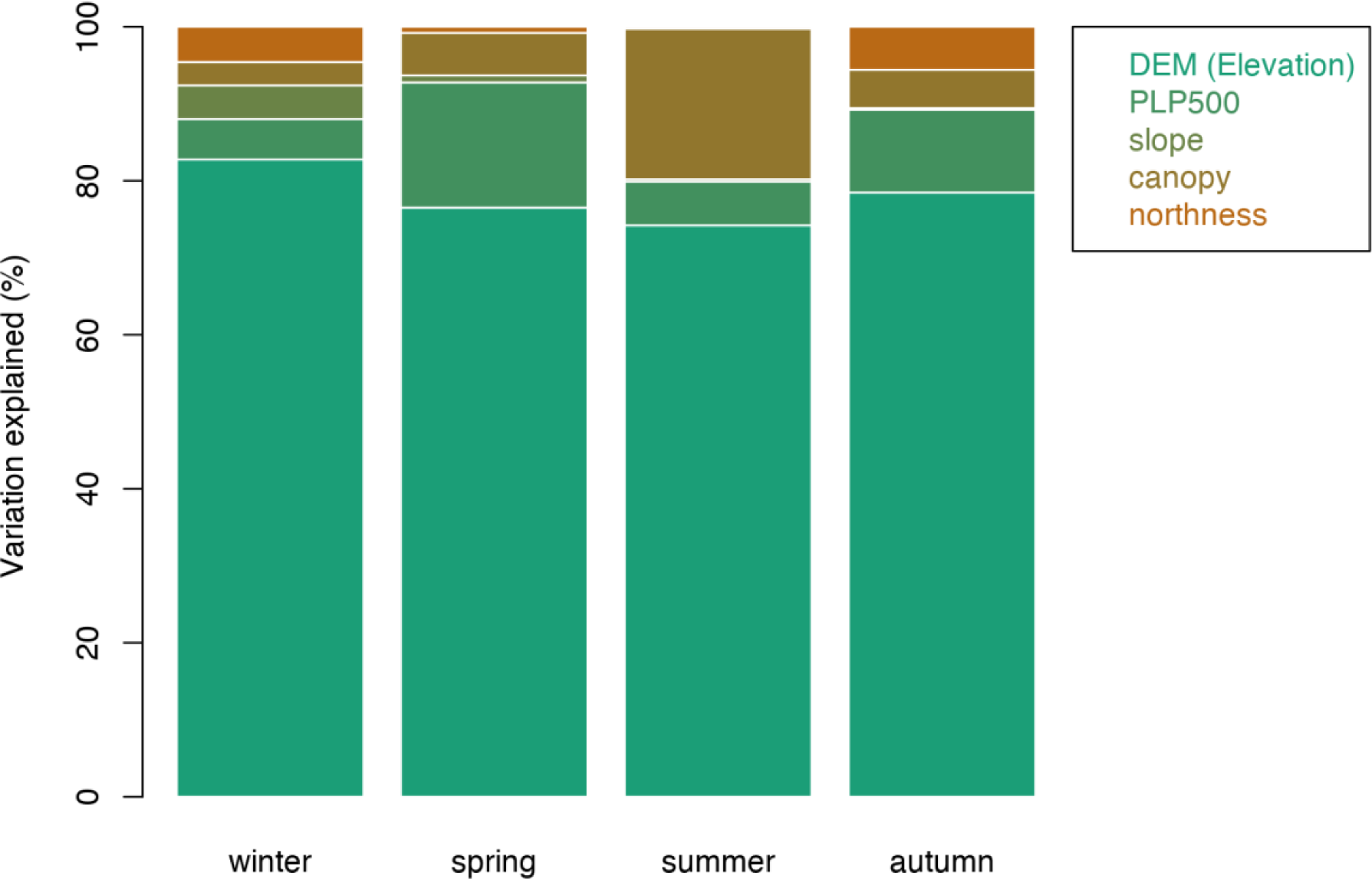
Topographic variables contributions deduced by RDA across all the seasons. Elevation was found to be a prominent driver of variability in all the seasons with contributions from other topographic features.

Inversion of minimum temperatures was found across Pepperwood Preserve in all the seasons. Lower elevation sites are colder than the higher elevation sites, especially in winter as evidenced by the steeper slopes (Fig. 6A). Autumn and spring are intermediate, with a slightly stronger pattern in autumn (Fig. 6A). Unlike the minima, maximum temperatures decline with elevation, and the pattern was weakest in winter (Fig. 6B). Higher elevation sites have lower diurnal variation in all seasons as opposed to lower elevation valley sites (Fig. 7 A). Across all the seasons, diurnal variations were highest in summer with the lowest experienced in winter.

**Figure 6:**
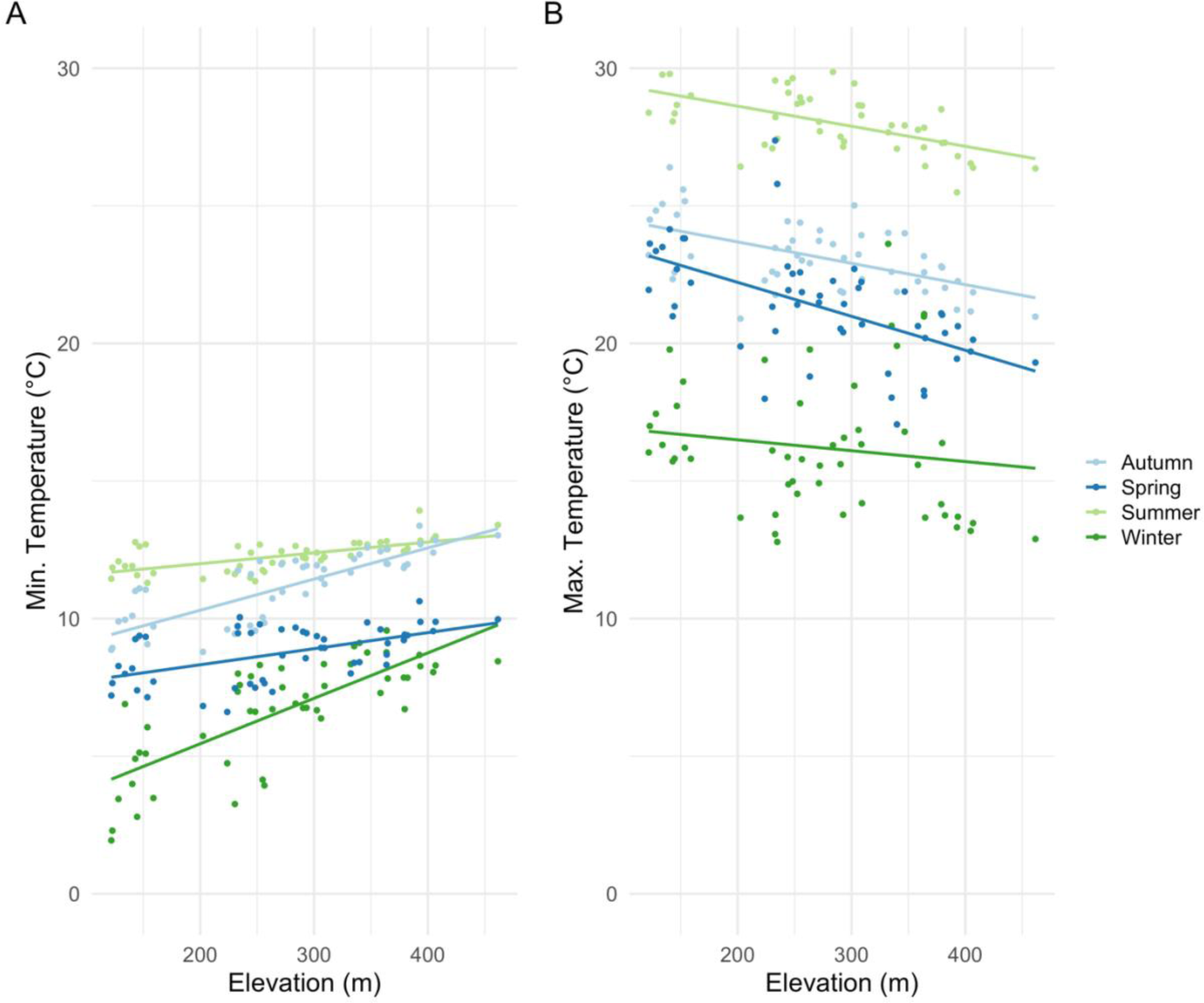
Linear fits of seasonal minima and maxima across the elevational gradient. (A) Lower elevation sites are found to be colder than the high elevation suggesting inversion (temperature increasing with elevation); autumn minimum temperatures exhibiting higher rate of increase by elevation than in the spring; Autumn (R^2^ = 65%, p-value < 0.001), Spring (R^2^ = 28%, p-value < 0.001), Summer (R^2^ = 41%, p-value < 0.001), and Winter (R^2^ = 58%, p-value < 0.001); (B) Summer maximum temperatures are found to be marginally varying with elevation; maximum temperatures in autumn are found to be less steep than spring; Autumn (R^2^ = 31%, p-value < 0.001), Spring (R^2^ = 31%, p-value < 0.001), Summer (R^2^ = 45%, p-value < 0.001), and Winter (R^2^ = 2%, p-value = 0.31).

**Figure 7:**
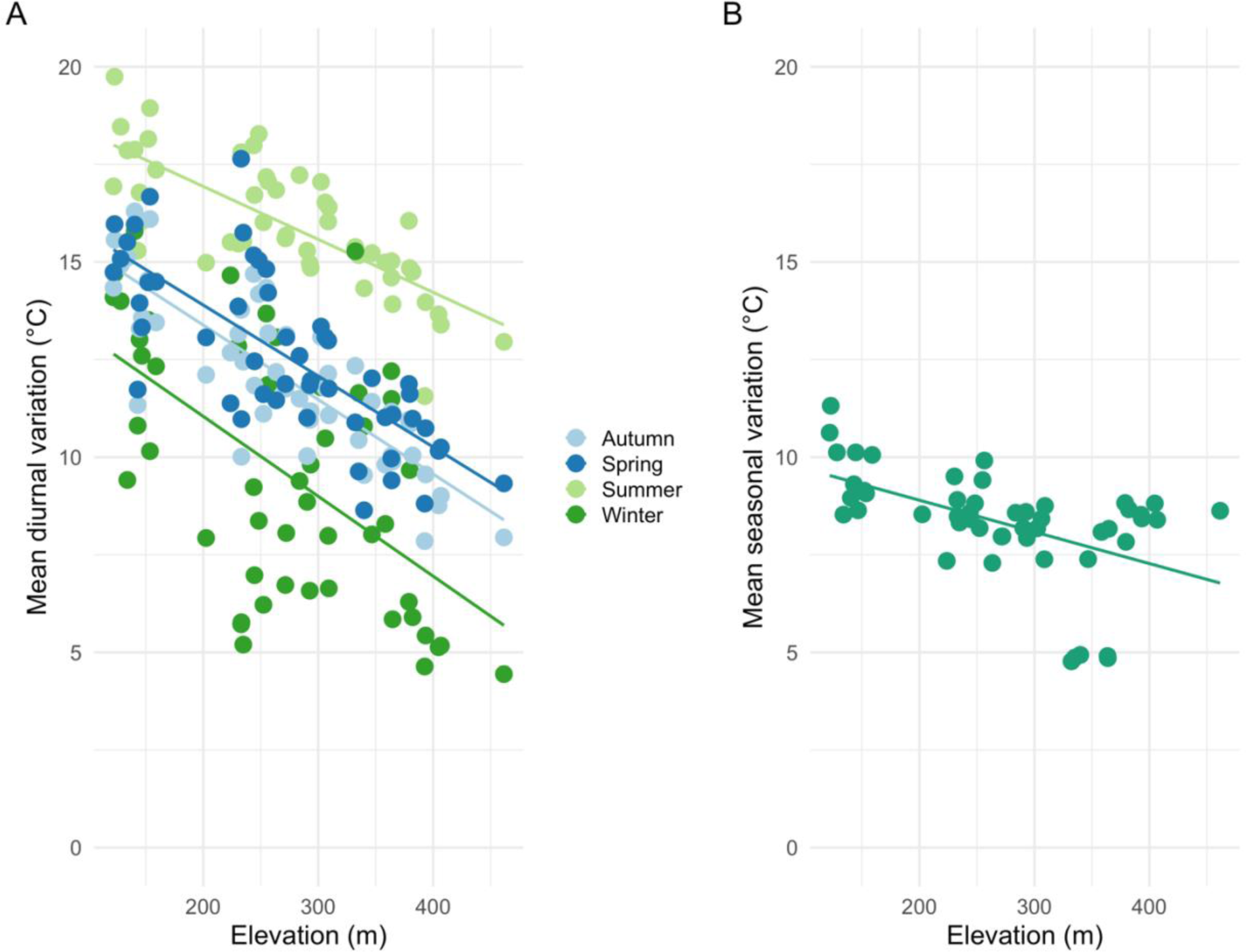
Linear fits of seasonal diurnal fluctuations as a function of elevation. Higher elevation sites exhibit lower diurnal variation in all seasons; Autumn (R^2^ = 69%, p-value < 0.001), Spring (R^2^ = 57%, p-value < 0.001), Summer (R^2^ = 57%, p-value < 0.001), and Winter (R^2^ = 31%, p-value < 0.001) (A) and lower seasonal variation; R^2^ = 25%, p-value < 0.001(B).

Comparison of daily mean temperatures of HOBOs with Pepperwood weather station showed a general trend of buffered temperatures in the understory than the weather station (Fig. 8, Fig. B.1). Majority of sites were found to be cooler in spring and summer, but warmer in fall and winter. The pattern was not evident in some of the high elevation sites (cluster of sites around 400 m elevation) where it tended to be nearly aligned with the weather station (Fig. 8B). Furthermore, comparison of diurnal variation of loggers with weather station shows lower diurnal variation of valley bottom sites than the high elevation sites with respect to weather station (Fig. B.2)

**Figure 8:**
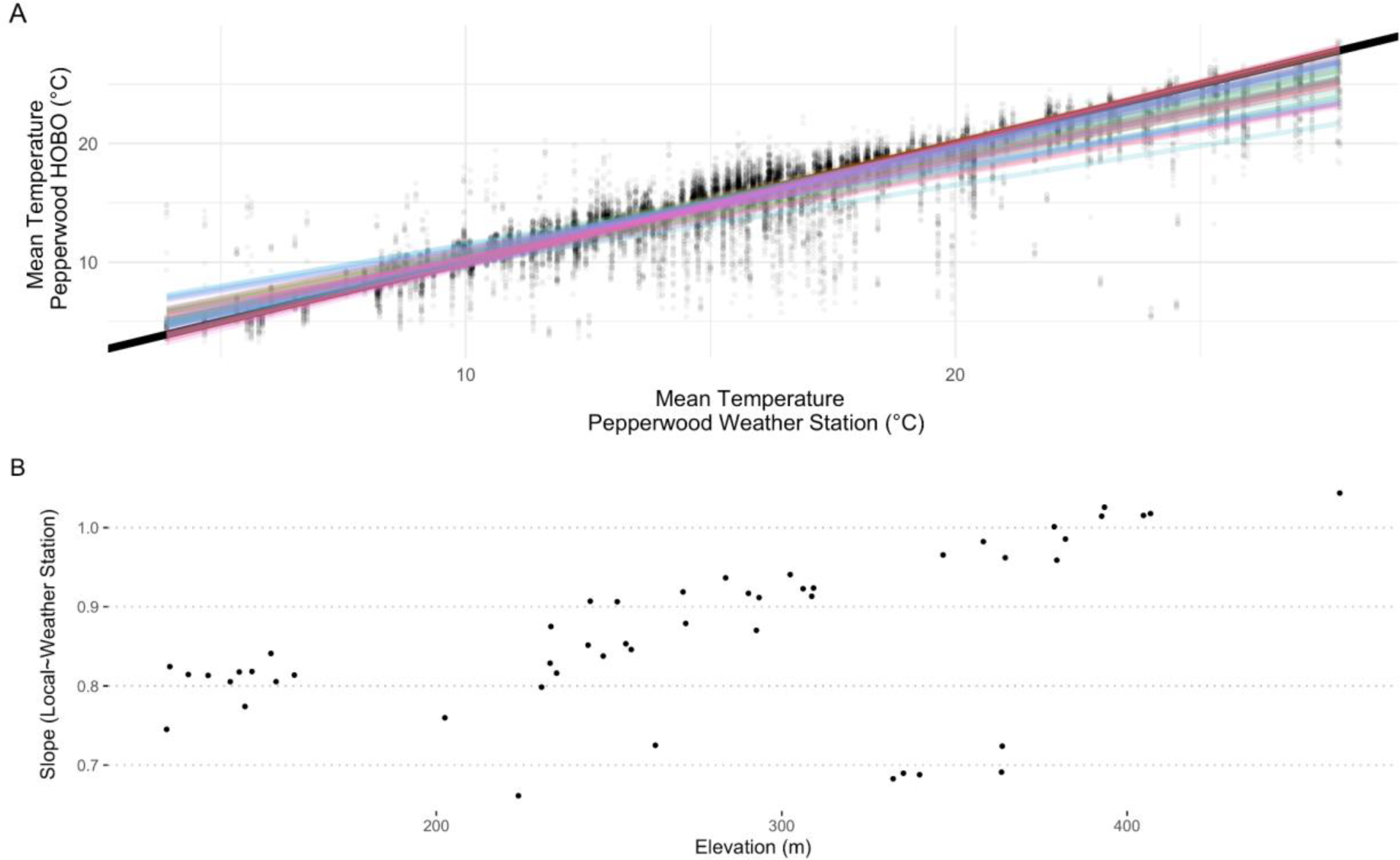
Linear fit of mean daily temperatures of HOBOs against the Pepperwood Preserve weather station (A) and slope of the relationship with elevation (B). Relationship highlighting that most of the sites are cooler in comparison to the weather station, but also show some of the high elevation sites are fundamentally aligned with the weather station climatically. Note that in panel A, dots refer to the day of the year, solid black line is 1:1, and the colored lines are individual linear fits in relation to the weather station.

## Discussion

Our study supports the general understanding that elevation is the primary driver for temperature variation, but also illustrates the additional influence of local vegetation and topography on the climatic conditions manifested at the site level in mountainous terrain. We confirm the role of elevation at a larger landscape level influencing both diurnal and seasonal temperature fluctuations (Pepin and Lundquist 2008, Dobrowski 2011), but found it to be inverted for minimum and maximum temperatures (Fig. 6A, B). Additionally, topographic features like topographic position (PLP500, direct measure of hilltop/valley bottoms) and northness contributed to the diurnal and seasonal variability, minima, and maxima. We also revealed that canopy mediates the diurnal variation in different seasons. Valley-bottom sites were in a cold-air pool and showed more variability (diurnal and seasonal), thus supporting our main hypothesis. Given that our results align with general understanding of elevation as the main contributor to temperature variation, it supports downscaling approaches that use coarse-grain climate grids to produce fine-grained products using only the elevation (Davy and Kusch 2021). However, one needs to be cautious as seasonal and region-specific lapse rates are to be considered when downscaling (Minder et al. 2010). Additionally, our study only used a year of climate data to look at topographic effects, a longer time period is warranted to ascertain the uncovered patterns.

Diurnal and seasonal temperature fluctuations are largely influenced by elevation but moderated by a combination of landscape and physiographic features. Average diurnal variations across the sites in all seasons were influenced by elevation (DEM) with lower elevations showing larger diurnal variation (Fig. 7). Furthermore, in our study site, we find that the more exposed a site is, like the ridgetops, the fluctuations are greater than the free-air temperature than the valley bottom sites that tend to be lower ((Dobrowski 2011), Fig. B.2). Diurnal fluctuation in different seasons were strongly influenced by topographic position and moderated by canopy, similar to findings in previous studies (Frey et al. 2016, Greiser et al. 2018, Davis et al. 2018, 2019).

Winter minimum temperatures were not influenced by finer physiographic features (Fig. 6A). However, ridgetops showed higher summer and winter mean temperatures than the valley bottoms. Our finding is consistent with past works that suggest valley bottoms receive less radiation load in comparison to ridgetops (Pepin and Lundquist 2008, Davis et al. 2019). Winter temperatures were found to be increasing with elevation (Fig. 6). The northwest portion of the Pepperwood Preserve exhibited strong inversion patterns, and potentially is part of a large-scale cold-air pool linked to a larger valley to the northwest of our study area (Fig. A.4).

In winter and autumn, we see that temperatures are driven by northness (Fig. 5), which agrees with past studies on the importance of aspect for radiation load. In winter, maximum temperatures were only explained by northness, reflecting effects of weaker intensity of solar insolation as angles are low. Also, spatially, winter max temperatures were more variable than any of the other seasons, likely suggesting cold-air pooling effects in winter. Less direct sun in winter as opposed to higher sun angles in summer likely explain the canopy effects that are found only in summer including seasonality in canopy cover (more effective shading by trees). A recent study at Pepperwood Preserve reported that site aspect was the most important determinant of species distributions of trees (Ackerly et al. 2020). But as our temperatures are from understories, mediated by canopy cover, it is most relevant to seedlings/understory vegetation rather than the overstory (Perry and Wu 1960). Our findings suggest possible stronger effects of winter minima (because of cold-air pools) that may influence future species distributions, more than summer maxima (Jackson et al. 2009).

We infer that the lower elevation valley sites that also happen to be in cold-air pools would be more unstable microclimatically (larger daily fluctuations) than the high elevation sites. In our study area, low elevation valley bottom sites are strongly buffered and high elevation sites are the most coupled (used weather station as a proxy to free-air; Fig. B.2, Fig. 7 and Fig. 8,). A variation indicative of the underlying topographic feature (Vanwalleghem and Meentemeyer 2009, Dobrowski 2011, McCullough et al. 2016, Zellweger et al. 2019). Physiographic features influence the degree of coupling of a site to the free-air temperature (Whiteman et al. 2004, Dobrowski 2011, Frey et al. 2016).

Canopy cover in conjunction with topographic attributes play a role in diurnal and seasonal fluctuations. We find the higher elevation sites that exhibit less diurnal and seasonal variation have lower canopy cover as opposed to lower valley bottom sites which exhibit higher variations (diurnal and seasonal). Canopy that has been noted in many studies to be contributing to microclimatic buffering is found here to be contributing to diurnal/seasonal variability (Greiser et al. 2018). One caveat to our study is that it is possible that we did not find very strong effects of canopy cover because all our loggers were under canopies (range of 30 - 90 %). However, as expected, the buffering effects of forest canopies are strongest when compared with an open site (De Frenne et al. 2019, 2021) (see Fig. 8). So, canopy cover mediates regional and macroclimatic climatic patterns in relation with elevational gradient and is likely contributing to lower variability diurnally and seasonally (Augusto et al. 2003). Also, despite having a short canopy cover gradient, we see strong effects of canopy in the summer (Fig. 5); this could be particularly important for management and conservation (Keppel et al. 2017).

### Implications for global change biology

The results presented here highlight the importance of microclimates, such as valley bottoms, that may be both cooler (at night and in winter) and warmer (during the day and in summer), compared to more exposed sites that are strongly coupled with regional climate. This pattern raises two sets of questions related to impacts of future climate change. First is the question of whether sites that are more variable diurnally or seasonally (i.e., valley bottoms) will change more or less in response to regional climate change (Maclean et al. 2017). The null expectation is that warming will be equivalent across sites, so the patterns of diurnal and seasonal variability will be maintained but all sites will be warmer. In midlatitudes it is widely observed that nighttime warming is greater than daytime (Yu et al. 2017, Slot and Winter 2018). Thus, diurnal variability should be declining across the landscape, though we are not aware of direct quantification of this effect. This change may lead to species distribution shifts at cold temperature limits, such as downslope movement into valleys as cold-air pools shrink (see (Van de Ven et al. 2007)), or (in coastal California) shifts away from the coast for species sensitive to winter frost. However, we are not aware of observations or first principles analysis addressing whether nighttime warming is greater in some sites vs. others, at a fine topographic scale.

The second set of questions is concerned with the impact of new extreme high temperatures on species (Vasseur et al. 2014). The comparison of hilltop to valley bottom sites is analogous to studies of maritime vs. continental climates (Ramirez et al. 2020), or tropical vs. temperate climates (Janzen 1967), in terms of the contrasts in diurnal and seasonal means and consequences for responses to climate change. By analogy there are two new contrasting hypotheses. The first is that the buffered sites with cooler daytime and summer high temperatures will remain cooler in the future, so they will be buffered from the most extreme impacts of climate change. Alternatively, the species occupying these sites may be adapted to the narrower range of temperatures, and thus be more sensitive to rising temperatures. By contrast, those that occupy sites with high diurnal and seasonal variability may also be better adapted to withstand new extremes under a changing climate (Helmuth et al. 2014).

## Limitations

We want to emphasize that the study was carried out on only one year of data, so one must be cautious on drawing broad scale inferences. Secondly, the study was carried out in a particular coastal landscape, so the inferences arrived here might not be the same if a similar study is carried out at a different landscape. However, our results confirm that elevation explains most of the seasonal variation in temperature. Next, we found the influence of cold-air pools in the observed temperatures in our study system, and one needs to be aware that such phenomena affect seasonal and region-specific lapse rates, and that might not be the case in a different study system (Curtis et al. 2014). Nonetheless, this study provides ecologists and global change biologists a way to interpret downscaled climate variables and the topographic, canopy, seasonally interactions they may be incorporating.

## Conclusion

Our study quantifies the importance of finer physiographic features like topographic position, northness and canopy on diurnal and seasonal variability. Also, we confirm the role of elevation and suggest that interactions with finer physiographic features would be key to understanding current and future species distributions. It is likely that sites having a larger diurnal or seasonal variation (valley sites in our study) might also be the same sites that are buffered from regional climatic patterns, and thus would most likely be better adapted to withstand new extremes under a changing climate. Though this study was limited to one-year, future work spanning multiple years can be done to ascertain buffering from climate change offered by temperature refugia (different rate of warming). Buffered areas can protect native species and ecosystems from the negative effects of climate change in the short term and provide longer-term havens from climate impacts for biodiversity and ecosystem function (Keppel et al. 2012, Morelli et al. 2020).

## Contributions by the Authors

A.J., and D.D.A. conceived and designed the research. A.J. performed the research. A.J. analyzed the data. A.J., J.D.O., M.F.O., M.M.K. and D.D.A. wrote the paper.

## Conflicts of Interest

None declared.

## Acknowledgements

The authors thank stewards of Pepperwood Preserve (PP) that aided in the establishment and monitoring of the plots. The authors also thank the Ackerly Lab for logistical support and fruitful discussions. Comments from two anonymous reviewers greatly improved the final manuscript. Funding for establishment of the research plot network was provided by the Moore Foundation. A.J. was supported by grants from the Department of Biology, University of Washington.

## Appendices

## Appendix A

**Figure A.1:**
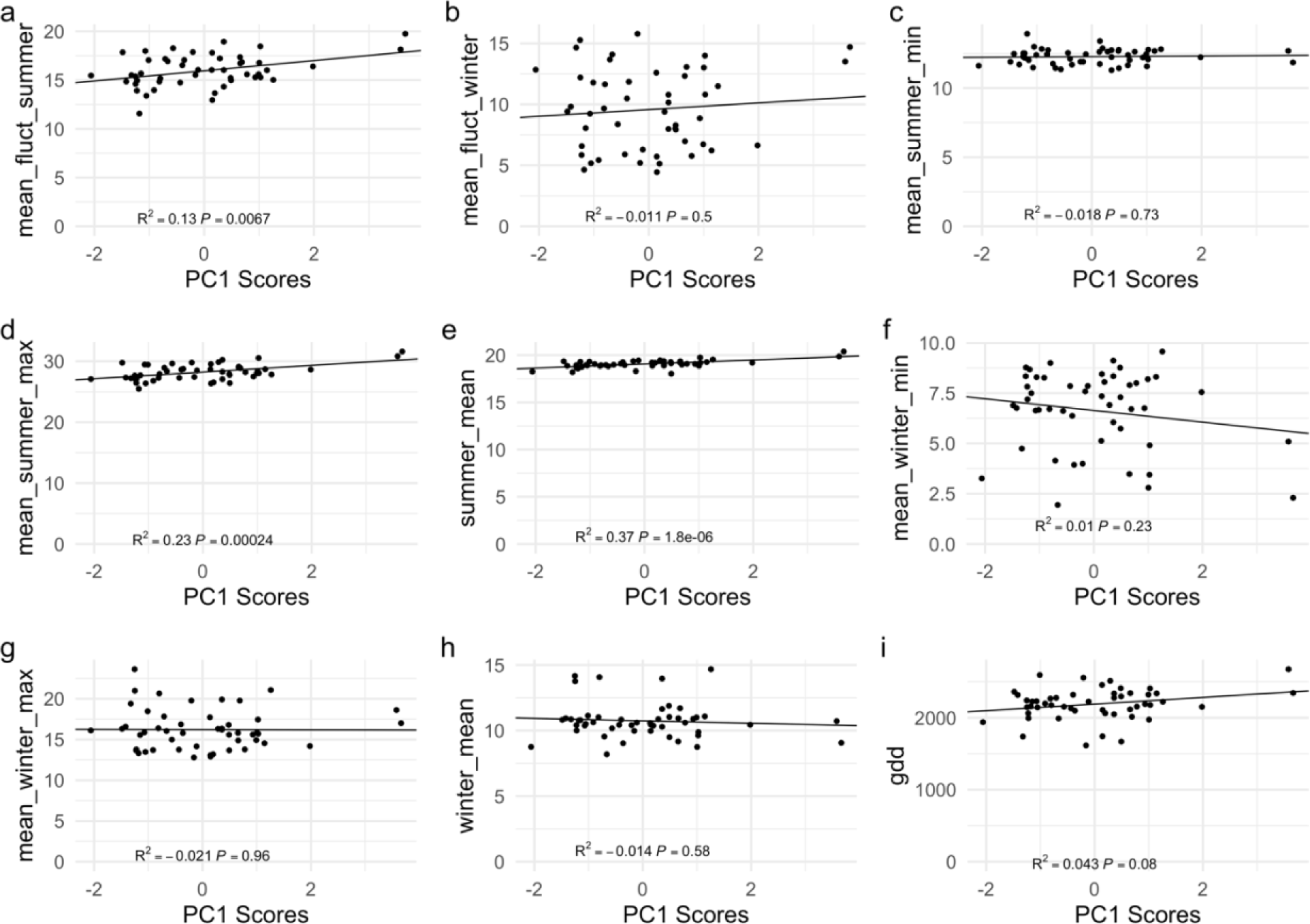
Regression of climate variables against PC1 of physiographic space. Dominant variable explaining PC1 is DEM (elevation)

**Figure A.2:**
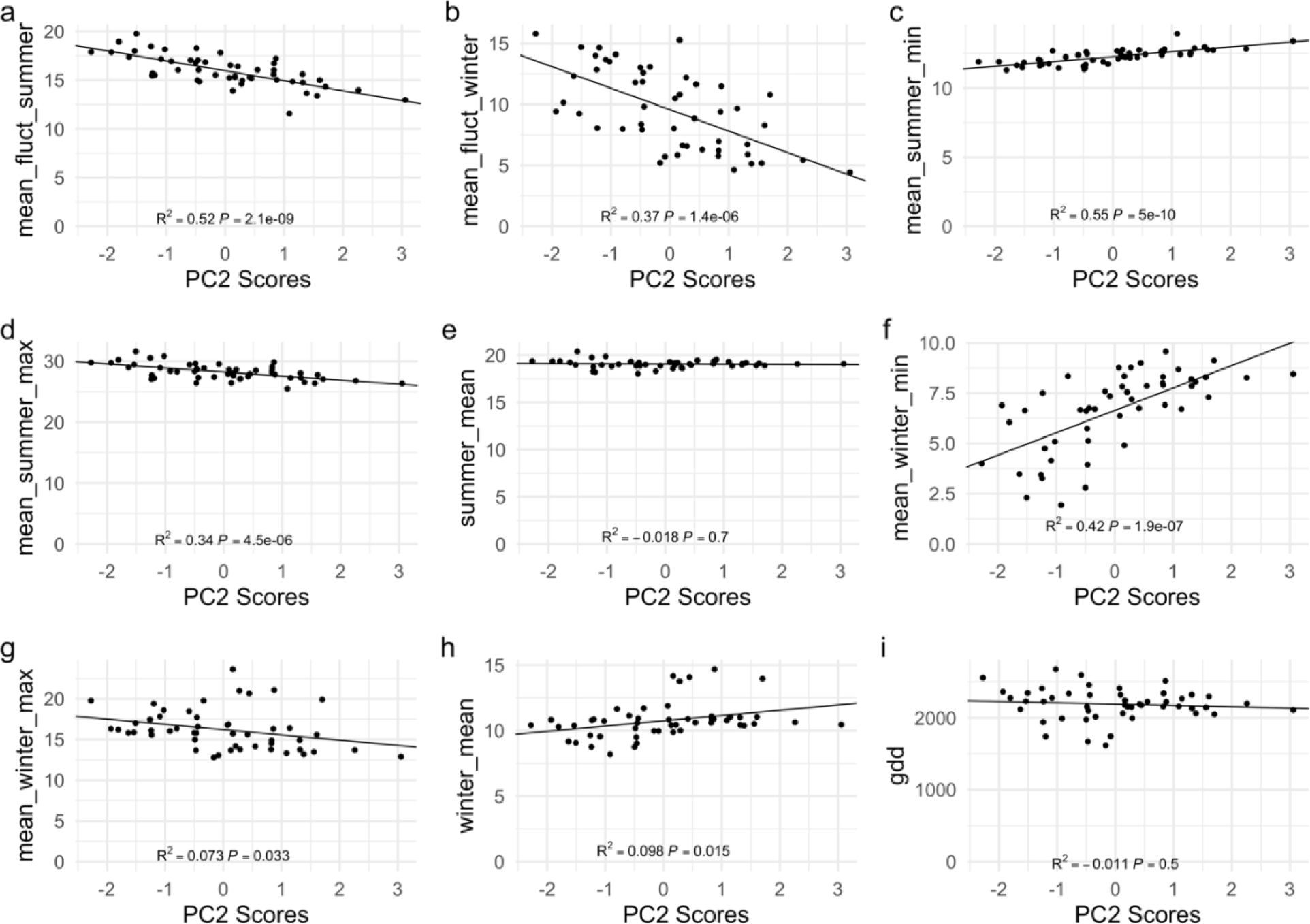
Regression of climate variables against PC2 of physiographic space. Dominant variables explaining PC2 are canopy, PLP500 and northness.

**Figure A.3:**
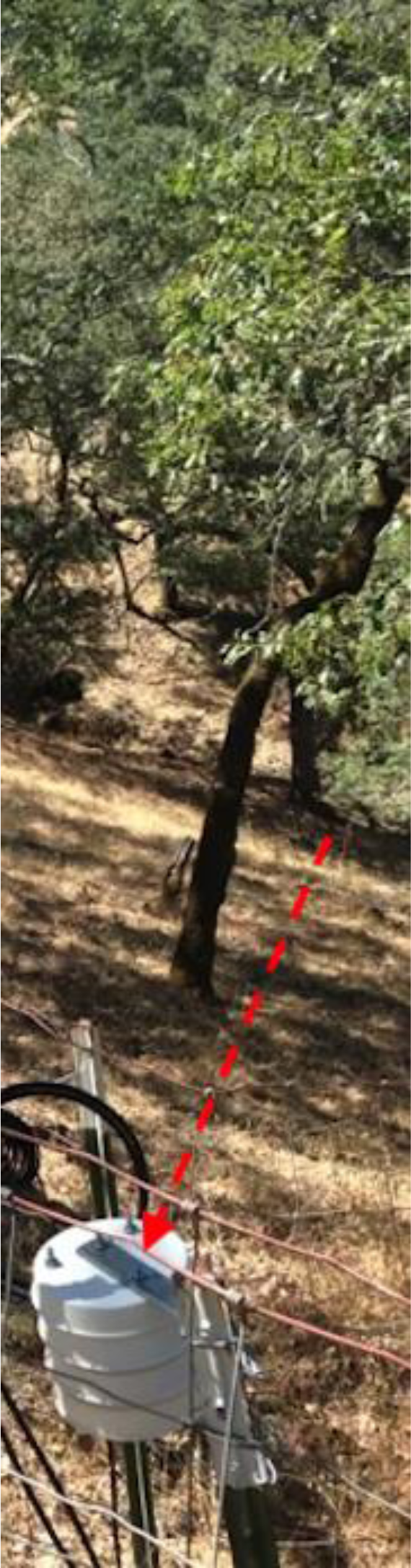
Custom radiation shield

**Figure A.4:**
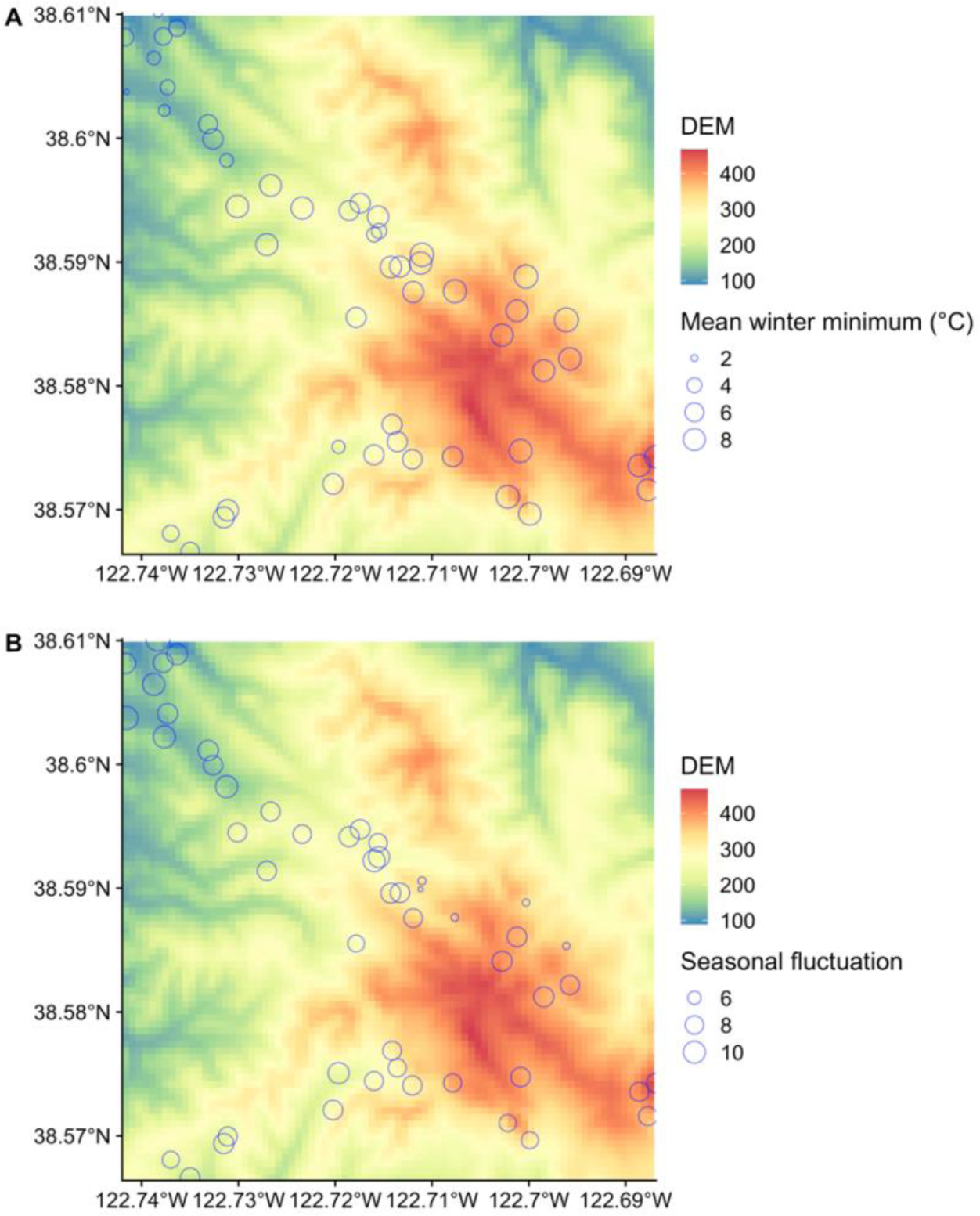
Cold-air pooling evident on the northwest corner Pepperwood Preserve. (a) Mean minimum temperatures highlight inversion. (b) Season fluctuation (summer mean – winter mean) pronounced in the upper northwest corner of the Pepperwood Preserve, possibly part of larger cold-air pooling phenomena.

**Table A.1.**
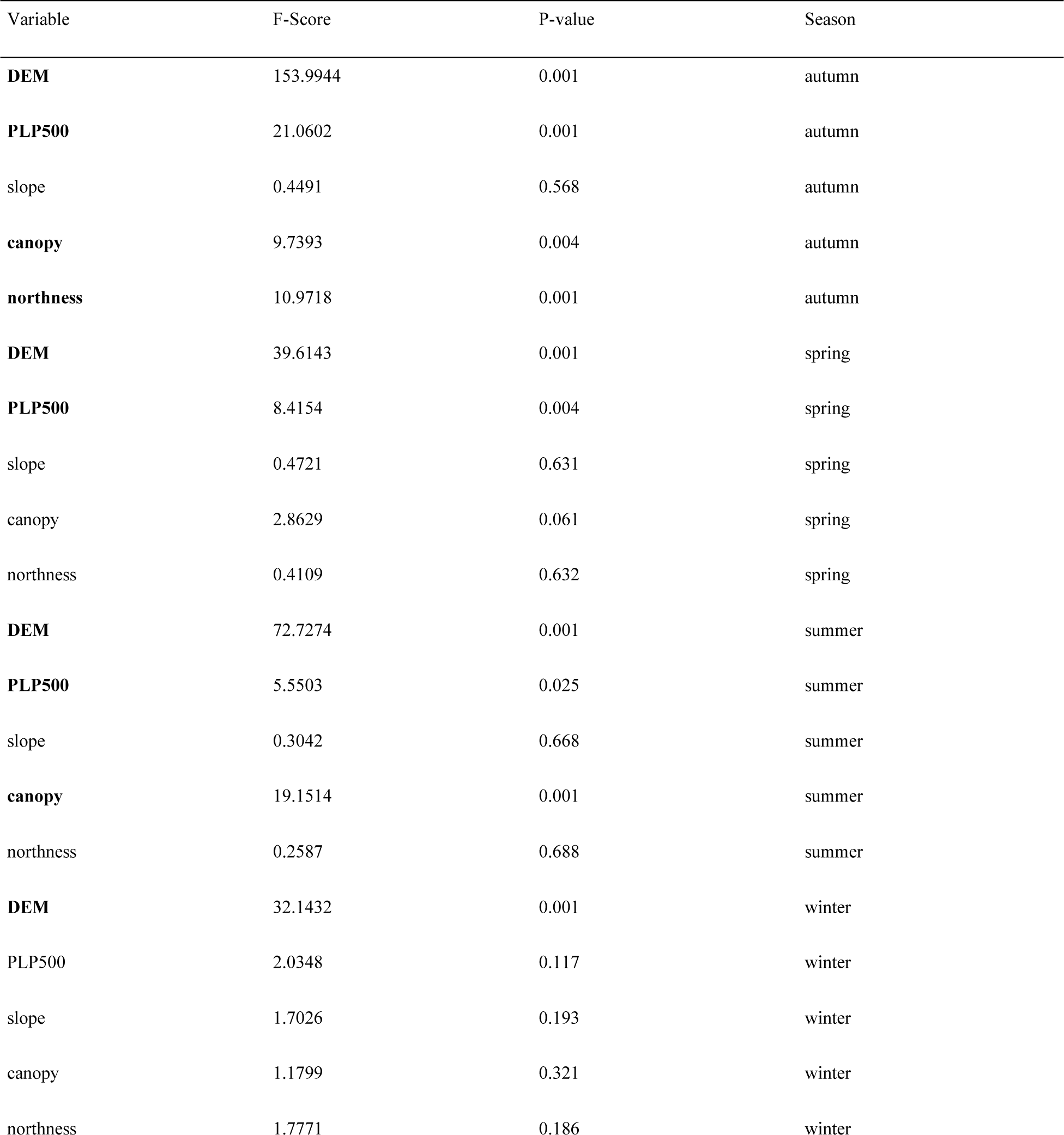
RDA results showing climate space (only seasonal metrics) when constrained against physiographic space. Variables in bold are statistically significant (p < 0.05).

**Table A.2.** Physiographic predictor variables and their effects.

| Variable | Effects |
| --- | --- |
| DEM (Elevation) | Temperature decreases with elevation |
| PLP500 | Topographic relief metric which provides indication of a valley bottom or a ridgetop. Valleys tend to be buffered from regional extremes whereas ridgetops generally follow the same pattern as regional climate. |
| slope | Influence solar radiation exposure (with aspect) |
| canopy | Canopies buffer daily temperature extremes |
| northness | Provides cooling effect during the day due to topography moderated solar radiation |

## Appendix B

**Figure B.1:**
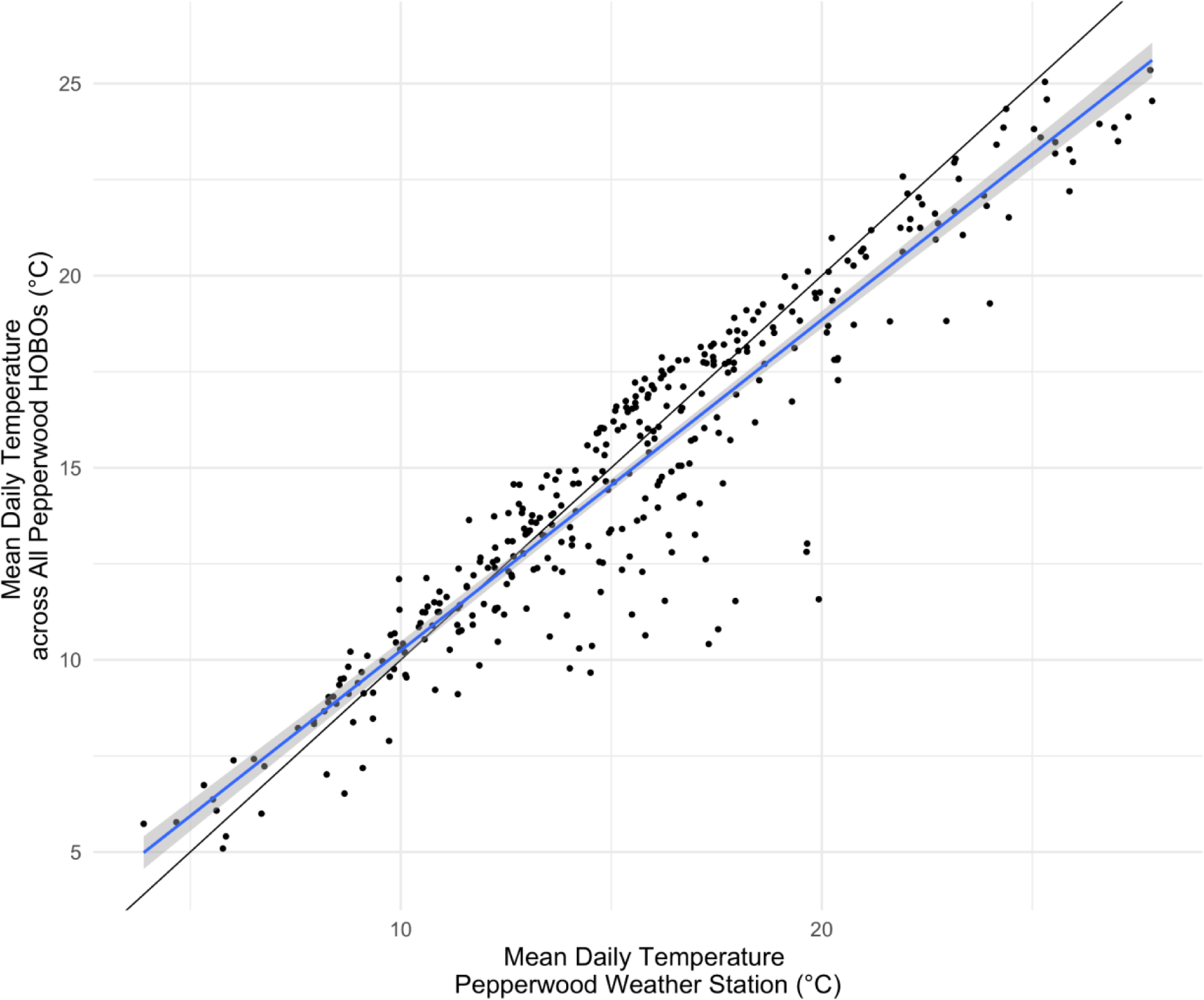
Mean temperature comparisons between HOBOs and an open site (Pepperwood Weather Station).

**Figure B.2:**
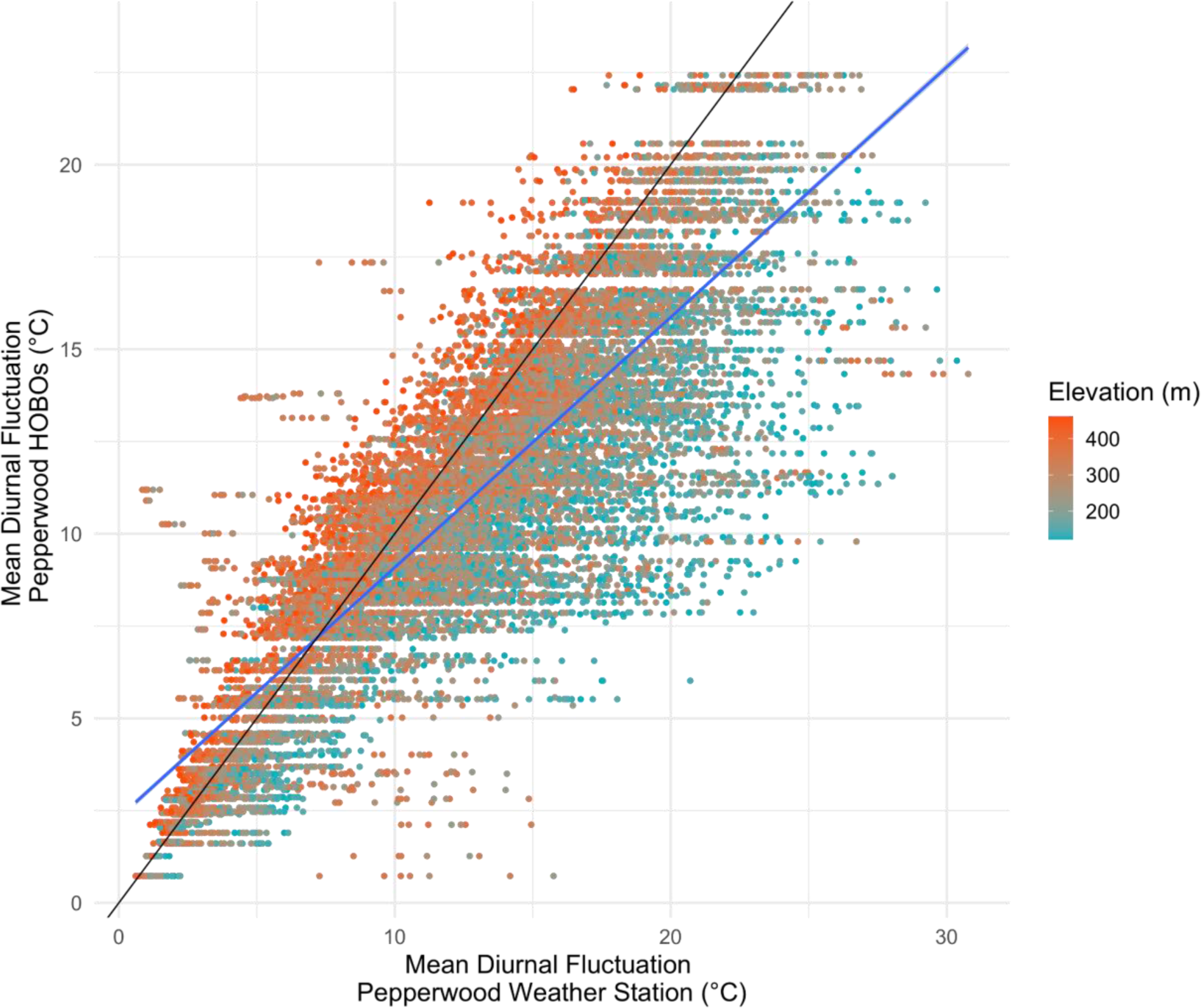
Mean diurnal fluctuation comparisons between HOBOs and an open site (Pepperwood Weather Station) colored by elevation.

## Notes

### Competing Interest Statement

The authors have declared no competing interest.

## References

Ackerly, D. D., M. M. Kling, M. L. Clark, P. Papper, M. F. Oldfather, A. L. Flint, and L. E. Flint. 2020. Topoclimates, refugia, and biotic responses to climate change. Frontiers in Ecology and the Environment 18:288–297.

Ashcroft, M. B., J. R. Gollan, D. I. Warton, and D. Ramp. 2012. A novel approach to quantify and locate potential microrefugia using topoclimate, climate stability, and isolation from the matrix. Global Change Biology 18:1866–1879.

Augusto, L., J.-L. Dupouey, and J. Ranger. 2003. Effects of tree species on understory vegetation and environmental conditions in temperate forests. Annals of Forest Science 60:823–831.

Burns, P., and C. Chemel. 2014. Evolution of Cold-Air-Pooling Processes in Complex Terrain. Boundary-Layer Meteorology 150:423–447.

Chen, I.-C., J. K. Hill, R. Ohlemuller, D. B. Roy, and C. D. Thomas. 2011. Rapid Range Shifts of Species Associated with High Levels of Climate Warming. Science 333:1024–1026.

Chen, J., and J. Franklin. 1997. Growing-season microclimate variability within an old-growth Douglas-fir forest. Climate Research 8:21–34.

Curtis, J. A., L. E. Flint, A. L. Flint, J. D. Lundquist, B. Hudgens, E. E. Boydston, and J. K. Young. 2014. Incorporating cold-air pooling into downscaled climate models increases potential refugia for snow-dependent species within the Sierra Nevada Ecoregion, CA. PLoS One 9:e106984.

Daly, C., D. R. Conklin, and M. H. Unsworth. 2009. Local atmospheric decoupling in complex topography alters climate change impacts. International Journal of Climatology:n/a-n/a.

Davis, K., S. Z. Dobrowski, Z. A. Holden, P. E. Higuera, and J. T. Abatzoglou. 2018. Microclimatic buffering in forests of the future: The role of local water balance. Ecography.

Davis, K. T., S. Z. Dobrowski, P. E. Higuera, Z. A. Holden, T. T. Veblen, M. T. Rother, S. A. Parks, A. Sala, and M. P. Maneta. 2019. Wildfires and climate change push low-elevation forests across a critical climate threshold for tree regeneration. Proceedings of the National Academy of Sciences 116:6193–6198.

Davy, R., and E. Kusch. 2021. Reconciling high resolution climate datasets using KrigR. Environmental Research Letters 16:124040.

De Frenne, P., J. Lenoir, M. Luoto, B. R. Scheffers, F. Zellweger, J. Aalto, M. B. Ashcroft, D. M. Christiansen, G. Decocq, K. De Pauw, S. Govaert, C. Greiser, E. Gril, A. Hampe, T. Jucker, D. H. Klinges, I. A. Koelemeijer, J. J. Lembrechts, R. Marrec, C. Meeussen, J. Ogée, V. Tyystjärvi, P. Vangansbeke, and K. Hylander. 2021. Forest microclimates and climate change: Importance, drivers and future research agenda. Global Change Biology 27:2279–2297.

De Frenne, P., F. Rodriguez-Sanchez, D. A. Coomes, L. Baeten, G. Verstraeten, M. Vellend, M. Bernhardt-Romermann, C. D. Brown, J. Brunet, J. Cornelis, G. M. Decocq, H. Dierschke, O. Eriksson, F. S. Gilliam, R. Hedl, T. Heinken, M. Hermy, P. Hommel, M. A. Jenkins, D. L. Kelly, K. J. Kirby, F. J. G. Mitchell, T. Naaf, M. Newman, G. Peterken, P. Petrik, J. Schultz, G. Sonnier, H. Van Calster, D. M. Waller, G.-R. Walther, P. S. White, K. D. Woods, M. Wulf, B. J. Graae, and K. Verheyen. 2013. Microclimate moderates plant responses to macroclimate warming. Proceedings of the National Academy of Sciences 110:18561–18565.

De Frenne, P., F. Zellweger, F. Rodríguez-Sánchez, B. R. Scheffers, K. Hylander, M. Luoto, M. Vellend, K. Verheyen, and J. Lenoir. 2019. Global buffering of temperatures under forest canopies. Nature Ecology & Evolution 3:744–749.

Dobrowski, S. Z. 2011. A climatic basis for microrefugia: The influence of terrain on climate. Global Change Biology 17:1022–1035.

Dorninger, M., C. D. Whiteman, B. Bica, S. Eisenbach, B. Pospichal, and R. Steinacker. 2011. Meteorological Events Affecting Cold-Air Pools in a Small Basin. Journal of Applied Meteorology and Climatology 50:2223–2234.

Ewers, R. M., and C. Banks-Leite. 2013. Fragmentation Impairs the Microclimate Buffering Effect of Tropical Forests. PLoS ONE 8:e58093.

Frey, S. J. K., A. S. Hadley, S. L. Johnson, M. Schulze, J. A. Jones, and M. G. Betts. 2016. Spatial models reveal the microclimatic buffering capacity of old-growth forests. Science Advances 2:e1501392.

Greiser, C., E. Meineri, M. Luoto, J. Ehrlén, and K. Hylander. 2018. Monthly microclimate models in a managed boreal forest landscape. Agricultural and Forest Meteorology 250– 251:147–158.

Hampe, A., and R. J. Petit. 2005. Conserving biodiversity under climate change: the rear edge matters: Rear edges and climate change. Ecology Letters 8:461–467.

Helmuth, B., B. D. Russell, S. D. Connell, Y. Dong, C. D. Harley, F. P. Lima, G. Sará, G. A. Williams, and N. Mieszkowska. 2014. Beyond long-term averages: making biological sense of a rapidly changing world. Climate Change Responses 1:6.

Jackson, S. T., J. L. Betancourt, R. K. Booth, and S. T. Gray. 2009. Ecology and the ratchet of events: Climate variability, niche dimensions, and species distributions. Proceedings of the National Academy of Sciences 106:19685–19692.

Janzen, D. H. 1967. Why Mountain Passes are Higher in the Tropics. The American Naturalist 101:233–249.

Jucker, T., S. R. Hardwick, S. Both, D. M. O. Elias, R. M. Ewers, D. T. Milodowski, T. Swinfield, and D. A. Coomes. 2018. Canopy structure and topography jointly constrain the microclimate of human-modified tropical landscapes. Global Change Biology 24:5243–5258.

Keppel, G., S. Anderson, C. Williams, S. Kleindorfer, and C. O’Connell. 2017. Microhabitats and canopy cover moderate high summer temperatures in a fragmented Mediterranean landscape. PLOS ONE 12:e0183106.

Keppel, G., K. P. Van Niel, G. W. Wardell-Johnson, C. J. Yates, M. Byrne, L. Mucina, A. G. T. Schut, S. D. Hopper, and S. E. Franklin. 2012. Refugia: identifying and understanding safe havens for biodiversity under climate change: Identifying and understanding refugia. Global Ecology and Biogeography 21:393–404.

Lareau, N. P., and J. D. Horel. 2015. Dynamically Induced Displacements of a Persistent Cold-Air Pool. Boundary-Layer Meteorology 154:291–316.

Lembrechts, J. J., I. Nijs, and J. Lenoir. 2019. Incorporating microclimate into species distribution models. Ecography 42:1267–1279.

Lenoir, J., B. J. Graae, P. A. Aarrestad, I. G. Alsos, W. S. Armbruster, G. Austrheim, C. Bergendorff, H. J. B. Birks, K. A. Br\a athen, J. Brunet, H. H. Bruun, C. J. Dahlberg, G. Decocq, M. Diekmann, M. Dynesius, R. Ejrn\a es, J. A. Grytnes, K. Hylander, K. Klanderud, M. Luoto, A. Milbau, M. Moora, B. Nygaard, A. Odland, V. T. Ravolainen, S. Reinhardt, S. M. Sandvik, F. H. Schei, J. D. M. Speed, L. U. Tveraabak, V. Vandvik, L. G. Velle, R. Virtanen, M. Zobel, and J. C. Svenning. 2013. Local temperatures inferred from plant communities suggest strong spatial buffering of climate warming across Northern Europe. Global Change Biology 19:1470–1481.

Lundquist, J. D., and D. R. Cayan. 2007. Surface temperature patterns in complex terrain: Daily variations and long-term change in the central Sierra Nevada, California. Journal of Geophysical Research 112.

Maclean, I. M. D., A. J. Suggitt, R. J. Wilson, J. P. Duffy, and J. J. Bennie. 2017. Fine-scale climate change: modelling spatial variation in biologically meaningful rates of warming. Global Change Biology 23:256–268.

McCullough, I. M., F. W. Davis, J. R. Dingman, L. E. Flint, A. L. Flint, J. M. Serra-Diaz, A. D. Syphard, M. A. Moritz, L. Hannah, and J. Franklin. 2016. High and dry: high elevations disproportionately exposed to regional climate change in Mediterranean-climate landscapes. Landscape Ecology 31:1063–1075.

Minder, J. R., P. W. Mote, and J. D. Lundquist. 2010. Surface temperature lapse rates over complex terrain: Lessons from the Cascade Mountains. Journal of Geophysical Research 115:D14122.

Morelli, T. L., C. W. Barrows, A. R. Ramirez, J. M. Cartwright, D. D. Ackerly, T. D. Eaves, J. L. Ebersole, M. A. Krawchuk, B. H. Letcher, M. F. Mahalovich, G. W. Meigs, J. L. Michalak, C. I. Millar, R. M. Quiñones, D. Stralberg, and J. H. Thorne. 2020. Climate-change refugia: biodiversity in the slow lane. Frontiers in Ecology and the Environment 18:228–234.

Oldfather, M. F., M. N. Britton, P. D. Papper, M. J. Koontz, M. M. Halbur, C. Dodge, A. L. Flint, L. E. Flint, and D. D. Ackerly. 2016. Effects of topoclimatic complexity on the composition of woody plant communities. AoB Plants 8:plw049.

Parmesan, C. 2006. Ecological and Evolutionary Responses to Recent Climate Change. Annual Review of Ecology, Evolution, and Systematics 37:637–669.

Patsiou, T. S., E. Conti, N. E. Zimmermann, S. Theodoridis, and C. F. Randin. 2014. Topo-climatic microrefugia explain the persistence of a rare endemic plant in the Alps during the last 21 millennia. Global Change Biology 20:2286–2300.

Pecl, G. T., M. B. Araújo, J. D. Bell, J. Blanchard, T. C. Bonebrake, I.-C. Chen, T. D. Clark, R. K. Colwell, F. Danielsen, B. Evengård, L. Falconi, S. Ferrier, S. Frusher, R. A. Garcia, R. B. Griffis, A. J. Hobday, C. Janion-Scheepers, M. A. Jarzyna, S. Jennings, J. Lenoir, H. I. Linnetved, V. Y. Martin, P. C. McCormack, J. McDonald, N. J. Mitchell, T. Mustonen, J. M. Pandolfi, N. Pettorelli, E. Popova, S. A. Robinson, B. R. Scheffers, J. D. Shaw, C. J. B. Sorte, J. M. Strugnell, J. M. Sunday, M.-N. Tuanmu, A. Vergés, C. Villanueva, T. Wernberg, E. Wapstra, and S. E. Williams. 2017. Biodiversity redistribution under climate change: Impacts on ecosystems and human well-being. Science 355:eaai9214.

Pepin, N. C., and J. D. Lundquist. 2008. Temperature trends at high elevations: Patterns across the globe. Geophysical Research Letters 35.

Perry, T. O., and W. C. Wu. 1960. Genetic Variation in the Winter Chilling Requirement for Date of Dormacy Break for Acer Rubrum. Ecology 41:790–794.

Quintero, I., and J. J. Wiens. 2013. Rates of projected climate change dramatically exceed past rates of climatic niche evolution among vertebrate species. Ecology Letters 16:1095– 1103.

R Core Team. 2021. R: A Language and Environment for Statistical Computing. R Foundation for Statistical Computing, Vienna, Austria.

Ramirez, A. R., M. E. De Guzman, T. E. Dawson, and D. D. Ackerly. 2020. Plant hydraulic traits reveal islands as refugia from worsening drought. Conservation Physiology 8:coz115.

Schörghofer, N., S. Businger, and M. Leopold. 2018. The Coldest Places in Hawaii: The Ice-Preserving Microclimates of High-Altitude Craters and Caves on Tropical Island Volcanoes. Bulletin of the American Meteorological Society 99:2313–2324.

Sexton, J. P., P. J. McIntyre, A. L. Angert, K. J. Rice, and others. 2009. Evolution and ecology of species range limits. Annual Review of Ecology, Evolution and Systematics 40:415–436.

Sheridan, P. F. 2019. Synoptic-flow interaction with valley cold-air pools and effects on cold-air pool persistence: Influence of valley size and atmospheric stability. Quarterly Journal of the Royal Meteorological Society 145:1636–1659.

Slot, M., and K. Winter. 2018. High tolerance of tropical sapling growth and gas exchange to moderate warming. Functional Ecology 32:599–611.

Thurman, L. L., B. A. Stein, E. A. Beever, W. Foden, S. R. Geange, N. Green, J. E. Gross, D. J. Lawrence, O. LeDee, J. D. Olden, L. M. Thompson, and B. E. Young. 2020. Persist in place or shift in space? Evaluating the adaptive capacity of species to climate change. Frontiers in Ecology and the Environment 18:520–528.

Van de Ven, C. M., S. B. Weiss, and W. G. Ernst. 2007. Plant Species Distributions under Present Conditions and Forecasted for Warmer Climates in an Arid Mountain Range. Earth Interactions 11:1–33.

Vanwalleghem, T., and R. K. Meentemeyer. 2009. Predicting Forest Microclimate in Heterogeneous Landscapes. Ecosystems 12:1158–1172.

Vasseur, D. A., J. P. DeLong, B. Gilbert, H. S. Greig, C. D. G. Harley, K. S. McCann, V. Savage, T. D. Tunney, and M. I. O’Connor. 2014. Increased temperature variation poses a greater risk to species than climate warming. Proceedings of the Royal Society B: Biological Sciences 281:20132612.

Whiteman, C. D., S. Eisenbach, B. Pospichal, and R. Steinacker. 2004. Comparison of Vertical Soundings and Sidewall Air Temperature Measurements in a Small Alpine Basin. Journal of Applied Meteorology 43:1635–1647.

Yu, M., Q. Gao, C. Gao, and C. Wang. 2017. Extent of Night Warming and Spatially Heterogeneous Cloudiness Differentiate Temporal Trend of Greenness in Mountainous Tropics in the New Century. Scientific Reports 7:41256.

Zellweger, F., P. De Frenne, J. Lenoir, D. Rocchini, and D. Coomes. 2019. Advances in Microclimate Ecology Arising from Remote Sensing. Trends in Ecology & Evolution 34:327–341.

